# Crosstalk between paralogous ComRS quorum-sensing systems diversifies the competence/bacteriocin networks of oral streptococci

**DOI:** 10.1101/2022.10.14.512260

**Authors:** Jinbei W. Li, Ryan M. Wyllie, Paul A. Jensen

**Affiliations:** Department of Bioengineering, University of Illinois at Urbana-Champaign, Urbana, IL, USA; Carl R. Woese Institute for Genomic Biology, University of Illinois at Urbana-Champaign, Urbana, IL, USA; Department of Microbiology, University of Illinois at Urbana-Champaign, Urbana, IL, USA

## Abstract

ComRS quorum-sensing systems regulate competence and bacteriocin production in the mutans streptococci that cause dental caries (tooth decay). We recently discovered a new class of ComRS (Type IV) in the oral pathogen *Streptococcus sobrinus*. We now show that every species of mutans streptococci has at least one Type IV ComRS system. *S. mutans, S. ratti, S. macacae*, and *S. ferus* have both a Type II and a Type IV ComRS system; *S. criceti* has one Type IV ComRS system; and *S. downei* and *S. sobrinus* have two Type IV ComRS systems. Gene deletion and reporter assays reveal that, in general, competence is regulated by one ComRS system, while a second ComRS system—when present—regulates bacteriocin production. However, crosstalk in species with two ComRS systems leads to a diverse set of competence/bacteriocin network structures. In *S. mutans*, for example, the Type IV ComRS can potentiate competence by kickstarting the well-studied Type II system, while the Type II system shows spillover activation of the Type IV system. Differences in ComR DNA binding sequences determine the specificity of the ComR paralogs, and sometimes a single base pair change can integrate the two systems. Our work reveals similarities and differences among the competence/bacteriocin networks in naturally competent streptococci, suggesting that paralogous regulation may provide evolutionary flexibility in genetic networks.

## 1 Introduction

Natural competence is a physiological state where microbial cells can actively acquire extracellular DNA. Competence promotes genetic exchange, genome repair, and nutrient acquisition[1]–[3] and is tightly regulated. In streptococci, regulation is achieved through quorum sensing systems that also control bacteriocin production, thus integrating these two processes [4], [5]. Competence systems like the ComCDE system in *Streptococcus pneumoniae* and the ComRS system in *S. mutans* are linked to growth, pathogenicity, social interactions, antibiotic resistance acquisition, and response to the human immune system [4]–[9].

The ComRS quorum-sensing system was discovered in *S. thremophilus* and *S. salivarius* as a regulator of natural competence [10]. In the canonical ComRS pathway (Figure 1A), ComR activates the alternative sigma factor ComX when stimulated with internalized Com*X*-*I*nducing *P*eptide (XIP) produced by extracellular processing of ComS [5]. Homologous ComRS systems were discovered in other streptococci, including the intensively studied ComRS pathway in *S. mutans* (Figure 1B) [11]. Adding synthetic XIP to cultures of streptococci can induce competence, providing a powerful tool for manipulating the genomes of the oral streptococci [12].

**Figure 1:**
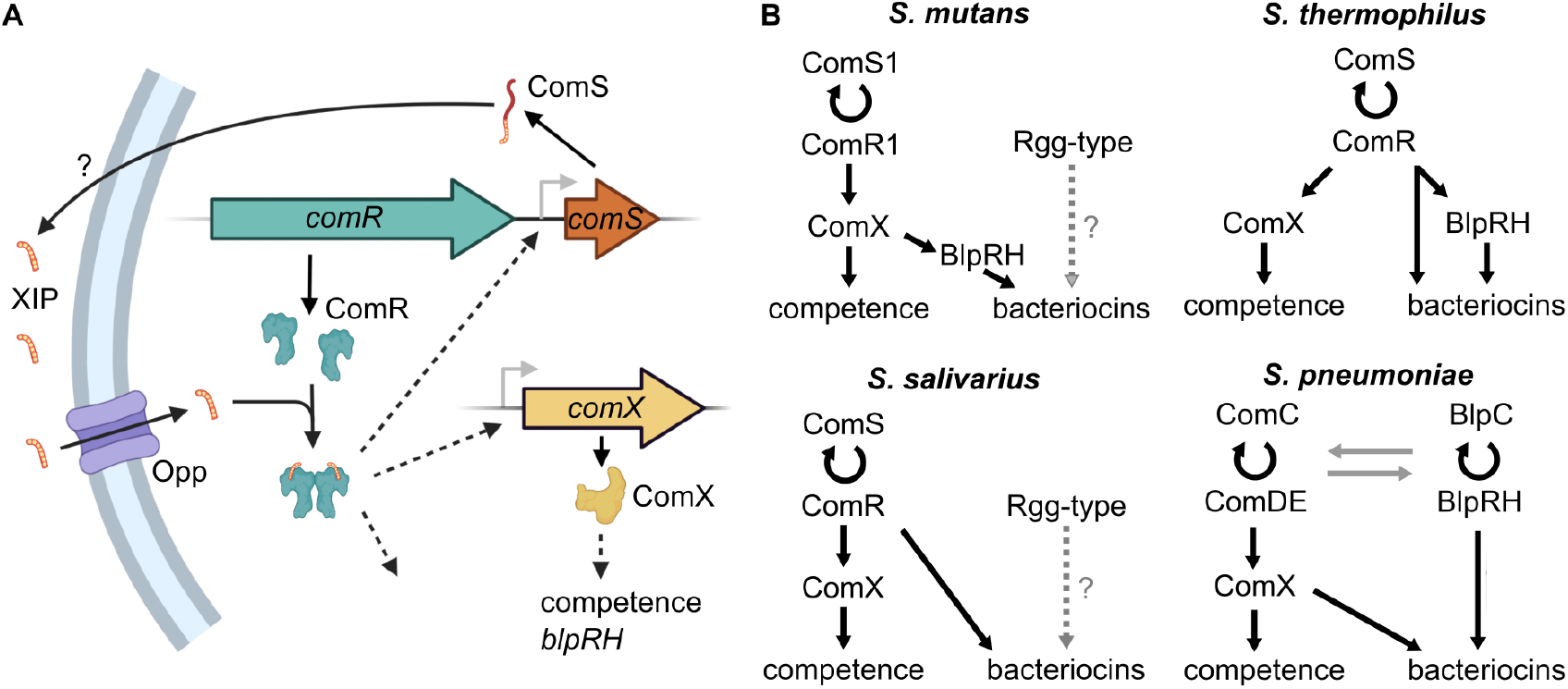
**A**. The ComRS pathway is an autocrine signaling loop. The ComS peptide is exported and cleaved into active XIP. XIP enters the cell and facilitates dimerization of ComR into an active transcription factor. **B**. Schematics of the competence-bacteriocin networks in different streptococci reflect knowledge before this study. In *S. mutans*, ComRS activates ComX, which subsequently activates competence genes and bacteriocins via BlpRH [13], [14]. *S. thermophilus* is similar, but ComR activates bacteriocins in the BlpRH system directly or through BlpCRH [15]. *S. salivarius*’s ComR activates ComX and directly activates bacteriocins [16]. A second ComR-like protein can also activate bacteriocins when given a synthetic peptide, but no ComS-encoding gene has been found for this pathway [17]. *S. pneumoniae* has dual two-component systems: ComCDE which controls competence and a group of bacteriocins, and BlpCRH which controls another group of bacteriocins; crosscontrol occur in both directions through transcriptional crosstalk and transporter promiscuity [18]–[20]. Dashed lines indicate hypothesized regulation. Gray arrows indicate indirect regulation.

Bacteriocins are peptides that lyse nearby cells or the producing cell. Coupling bacteriocins with competence may defend against other species [16], [21], lyse kin-cells to make homologous DNA available [22], or enhance quorum-sensing signals [23]. While competence genes are always under the control of ComX, which is in turn controlled by ComRS, the control of bacteriocin production varies by species. In *S. mutans*, the BlpCRH system that activates bacteriocin clusters is under control of ComX [13], [14]. In *S. thermophilus*, ComRS regulates bacteriocin genes in the BlpCRH system (a homolog of the ComCDE system) either directly or indirectly [15]. In *S. salivarius*, bacteriocin production is directly controlled by ComRS without either ComX or BlpCRH as intermediates [16]. In *S. pneumoniae*, a two-component ComCDE system (using CSP derived from ComC as an extracellular quorum-sensing signal) controls both competence and bacteriocins, while a paralogous BlpCRH system controls another group of bacteriocins; crosstalk in both directions exist between the ComCDE and BlpCRH system at the transcriptional level and through transporter promiscuity [18]–[20] (Figure 1B).

In 2020, we discovered a new class of ComRS system in *S. sobrinus* that differed from known ComRS systems in two ways [21]. First, the *S. sobrinus* XIP sequence lacks aromatic amino acids. All previously known XIPs contain two aromatic residues that are used to classify the respective ComRS systems: P(Y/F)F = Type I (salivarius group), WW = Type II (mutans, pyogenes, bovis, and agalactiae groups), and WG(T/K)W = Type III (*S. suis*) [5]. The *S. sobrinus* XIP implies that aromatic amino acids cannot reliably identify or classify ComRS systems. Instead, we now define a Type IV ComRS system as one with 1.) a XIP that does not fit into Types I–III, and 2.) a ComR that is more similar to the *S. sobrinus* ComR than to ComR proteins in the other classes.

The second novel feature of the *S. sobrinus* ComRS pathway is having two ComR proteins that function independently yet are associated with the same XIP. One protein (ComR2) is required for competence, and the presence of the other (ComR1) appears to inhibit competence [21]. *S. mutans* also contains a second putative ComR, but a corresponding ComS was never found and its function is unknown beyond a hypothesis that it regulates an adjacent bacteriocin cluster [11].

Here we show that all mutans streptococci contain at least one Type IV ComRS system. The previously reported second ComR in *S. mutans* is, in fact, a Type IV system with its own XIP. *S. ratti, S. macacae*, and *S.ferus* similarly contain both a Type II and a Type IV ComRS. *S. downei*, like *S. sobrinus*, contains two Type IV ComRS systems, but unlike *S. sobrinus* the *S. downei* ComRS systems have separate pheromones. *S. criceti* contains a single, Type IV ComRS system.

We also show that the ComRS-regulated network structures vary widely among mutans streptococci, leading to different levels of connectivity between competence and bacteriocin production. In species with both a Type II and Type IV ComRS system, competence is primarily controlled by the Type II pathway that sometimes regulates bacteriocins, while the Type IV pathway activates bacteriocins but does not directly regulate competence. In species with two Type IV systems, one system activates competence while the other controls bacteriocins. However, varying degrees of crosstalk exists between paralogous ComRS systems. For example, the two Type IV systems in *S. sobrinus* share a single XIP. The Type IV system in *S. mutans* can induce low levels of competence by kickstarting the Type II system, and the Type II system activates the Type IV system during peak of Type II expression. In *S. mutans*, we show that the specificity between the two ComRS systems is determined by a few base pairs in the ComR recognition sequences (ComR boxes) of the target promoters. In one case, changing a single nucleotide can integrate the Type II and Type IV ComRS systems. These results indicate high evolutionary flexibility and adaptability in species with dual ComRS systems.

**Table 1:**
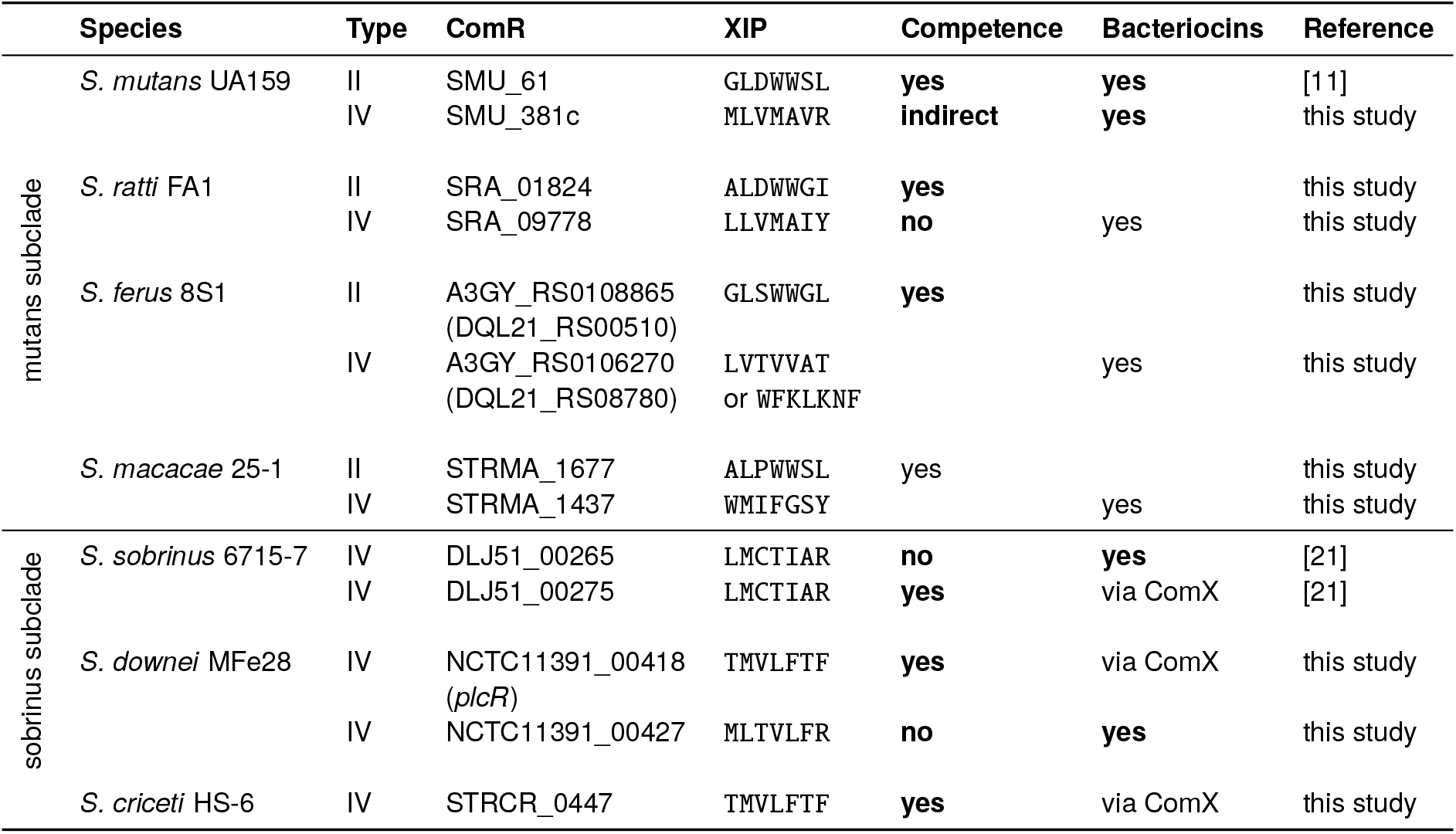
The mutans streptococci contain multiple ComRS systems that control competence and bacteriocin production. A bacteriocin (RiPP) gene cluster is predicted to be be in the ComRS network based on the presence of a ComR box in promoter region; bacteriocins nearby a *comRS* cluster are predicted to be controlled by it. Bold entries in these columns were experimentally validated by transformation or fluorescent reporter assays. For all three species in the sobrinus clade, a bacteriocin gene cluster is found to have a cin-box indicating ComX control, but induction is not observed in reporter assays.

## 2 Results

### 2.1 Mutans group streptococci have multiple ComRS systems

We searched for ComR homologs in the genomes of seven mutans streptococci based on the known ComR proteins in *S. mutans, S. sobrinus*, and *S. thermophilus* (Table 1). After identifying a *comR* gene, we searched nearby for a potential *comS* gene using the conserved promoter motif recognized by the ComR-XIP complex (the “ComR box”). The known Type II ComR in *S. mutans* has one homolog in each of three species: *S. ratti, S.ferus*, and *S. macacae*. The three putative *comR* genes all have a nearby *comS* encoding a peptide with the double-tryptophan motif that defines the Type II class [5]. We found no other Type II ComRS systems in the remaining mutans streptococci.

*S. sobrinus* has two *comR* genes (*comR1* and *comR2*) located in one gene cluster with a single Type IV *comS* following *comR1* [21]. As we show below, both ComR1 and ComR2 respond to the Type IV XIP, so we used both proteins to search for other Type IV ComRS systems in the mutans streptococci. All seven mutans streptococci, including the four with a Type II ComRS mentioned above (*S. mutans*, *S. ratti*, *S. ferus*, and *S. macacae*), contain at least one Type IV ComR with an adjacent ComS. Some of the predicted XIP sequences contain aromatic amino acids, but none match the motifs that define ComRS Types I, II, or III. The Type IV ComR in *S. mutans* (SMU.381c) was the aforementioned ComR with a “missing” ComS/XIP [11]. The Type IV *comS* gene we discovered is adjacent to SMU.381c, but it lacks the aromatic amino acids used to identify XIPs when the original *S. mutans* ComRS was discovered [11]. Two species, like *S. sobrinus*, have only Type IV ComRS systems—*S. criceti* has one Type IV ComRS, and *S. downei* has two Type IV systems, each with their own ComS.

### 2.2 ComRS classes follow the revised phylogeny of the mutans groups

The mutans streptococci were defined as a clade of cariogenic species containing the namesake *S. mutans*[24]. Recent phylogenetic reconstructions separate the mutans group into two subclades: a mutans subclade containing *S. mutans*, *S. ferus*, *S. ratti*, and *S. macacae*, and a new sobrinus subclade containing *S. sobrinus*, *S. criceti*, and *S. downei* [25], [26]. The distribution of ComRS systems follows the mutans/sobrinus split: all species in the new mutans subclade contain a Type II and a Type IV ComRS system, while species in the sobrinus subclade contain only Type IV ComRS systems (Table 1). Our original report of the Type IV ComRS in *S. sobrinus* noted that the ComR protein appeared more similar to Type I ComR proteins in the salivarius group [21], which is consistent with the placement of the sobrinus subclade nearer to the salivarius subclade than the mutans subclade [25], [26] (Figure 2).

**Figure 2:**
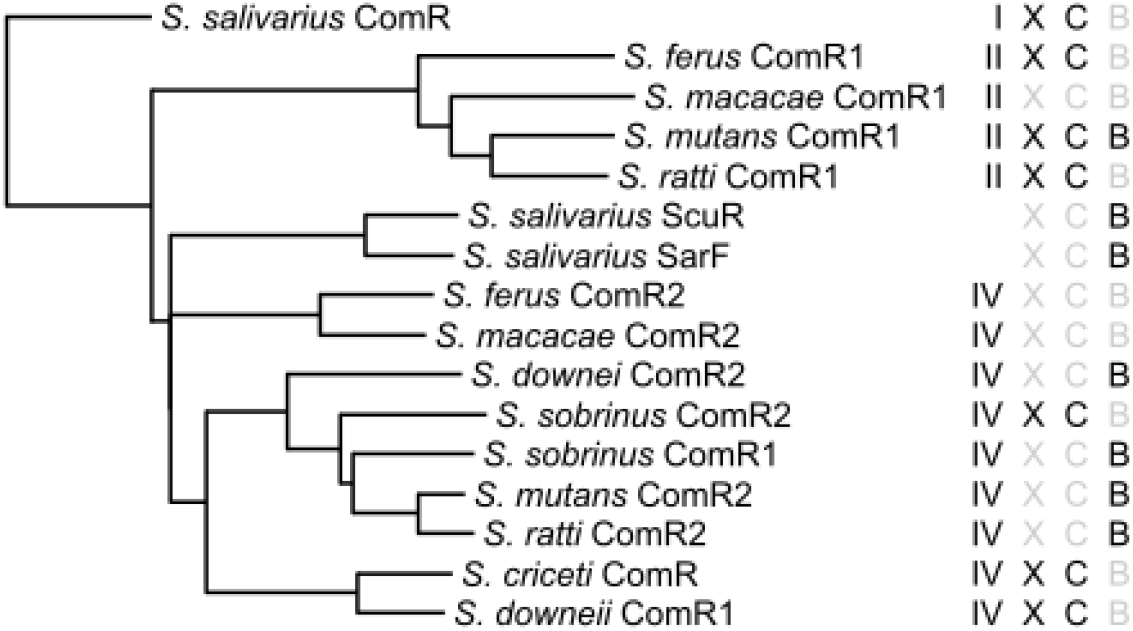
Streptococcal ComR sequences cluster according to class (Type II or Type IV) and follow the mutans/sobrinus subclades [25], [26]. Distances were computed using ClustalOmega, and each sequence is labeled by its class (I, II, or IV), if it activates ComX (X), induces competence (C), or upregulates bacteriocins (B). The recently discovered *S. salivarius* ScuR and SarF proteins lack a known pheromone but activate bacteriocins when stimulated with synthetic peptides not found in the genome [17]. *S. ratti* ComR2 is predicted to regulate an adjacent bacteriocin cluster, but we did not experimentally verify this prediction. We were unable to transform *S. macacae*, so the functions of its ComRS systems remain unknown.

### 2.3 Competence is predominantly activated by one ComRS system per species

In previously studied ComRS systems, the peptide XIP binds to ComR to upregulate the sigma factor ComX, which in turn activates natural competence [10], [11]. We tested if synthetic peptides based on the predicted XIP sequences could induce competence *in vitro* via transformation assays. Indeed, the *S. criceti* XIP induces competence and requires both the Type IV ComRS pathway and ComX (Figure 3A). In *S. downei*, we found two Type IV ComRS pathways, each with their own XIP. Only XIP1 (which is identical to the *S. criceti* XIP) induces competence and depends on the associated ComRS1 and ComX (Figure 3B). Deleting the ComRS2 pathway in *S. downei* doubles the transformation efficiency of XIP1 by one-fold (*p* = 0.02, *t*-test), suggesting that ComRS2 may inhibit competence (Figure 3B). This is similar to *S. sobrinus* where ComR2 is required for competence while ComR1 inhibits it [21].

**Figure 3:**
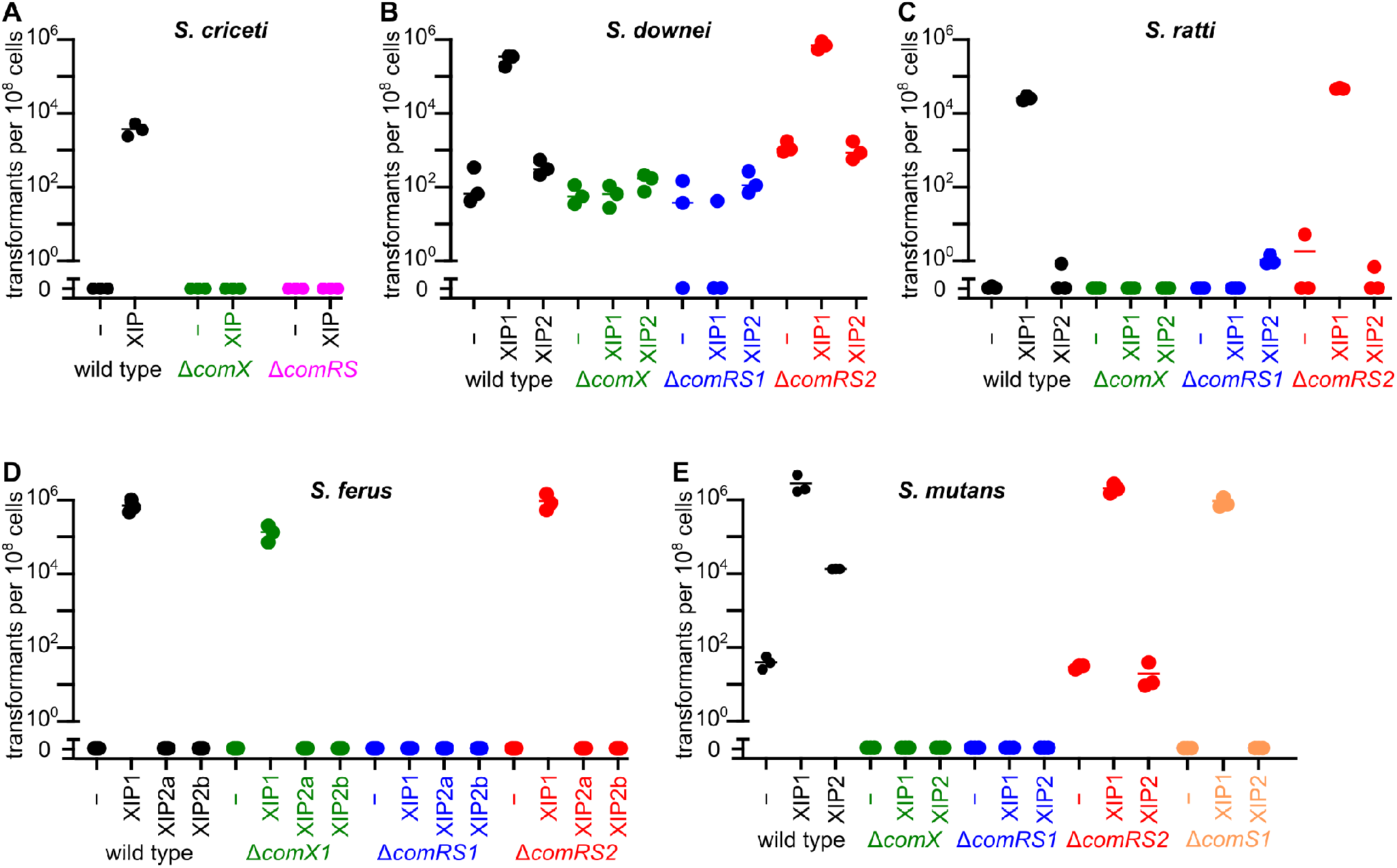
Competence is predominantly activated by one ComRS system in *S. criceti* (**A**), *S. downei* (**B**), *S. ratti* (**C**), *S. ferus* (**D**), and *S. mutans* (**E**). We previously showed that only one of the two ComRS systems in *S. sobrinus* controls competence [21]. **D**. *S. ferus* has two predicted XIP2 peptides (XIP2a and XIP2b) and two *comX* genes. Neither XIP2 induces competence. The *comX1* gene is not required for competence, although its deletion leads to 80% decrease in transformation efficiency. The well-known Type II ComRS systems in *S. mutans* is labeled ComRS1 to distinguish it from the Type IV ComRS2. The conditions use DMSO as a negative control. The ordinate “0” indicates transformants were not detected. Horizontal lines show the mean of three biological replicates.

*S. ratti* has both a Type II system (ComRS1/XIP1) and a Type IV system (ComRS2/XIP2). Exogenous XIP1 induces competence through the ComRS1 pathway and depends on ComX (Figure 3C). XIP2 never induced competence beyond background levels and deleting the ComRS2 pathway did not affect transformation efficiency (Figure 3C).

*S. ferus* similarly has a Type II system (ComRS1/XIP1) that induces competence and requires the *comRS1* genes. *S. ferus* also has a Type IV system with a single *comR2* gene and two potential *comS2* genes (*comS2a*/XIP2a and *comS2b*/XIP2b). Both *comS2* genes have a ComR box in their promoters, but the predicted XIP2a sequence has no aromatic amino acids (LVTVVAT) while the XIP2b sequence has three (WFKLKNF). Neither XIP2a nor XIP2b induces competence, and deleting the ComRS2 pathway does not affect the efficiency of transformation using XIP1 (Figure 3D). *S.ferus* is the only mutans streptococcus with two identical copies of the *comX* gene. Deleting one ComX reduced the transformation efficiency by 80% (Figure 3D).

We refer to the well-characterized *S. mutans* Type II ComRS system as ComRS1 to distinguish it from the newfound Type IV system ComRS2. It is well known that XIP1 induces competence through ComRS1 by activating ComX [11] (Figure 3E). The original paper describing ComRS1 also identified ComR2 (but not ComS2) and predicted that ComR2 could regulate bacteriocin production but not competence [11]. Surprisingly, the Type IV XIP2 we discovered also induces competence in *S. mutans* and requires both ComRS2 and ComX (Figure 3E). XIP2-induced competence also requires ComRS1 since a *ΔcomRS1* strain produced no transformants with either XIP1 or XIP2. A *ΔcomS1* strain can be transformed using XIP1 but not with XIP2, suggesting that XIP2 requires the positive feedback loop created by the autocrine ComR1/ComS1/XIP1 pathway (Figure 3E).

*S. macacae* contains a Type II and a Type IV ComRS—the former we predict controls competence. We were unable to transform *S. macacae* using either XIP1, XIP2, or XIP1 with one additional N-terminal amino acid (`RALPWWSL`). *S. macacae* remains the only genetically intractable species in the mutans group, and we necessarily excluded it from the genetic reporter assays below.

A pattern emerges from our transformation results. In transformable mutans streptococci with both a Type II and Type IV ComRS (*S. mutans, S. ratti*, and *S.ferus*), competence is controlled by the Type II ComRS but not the Type IV ComRS. The lone exception is *S. mutans*, where the Type IV XIP2 can induce low levels of competence but requires both the Type II and Type IV ComRS systems and ComX. In species with only Type IV ComRS systems (*S. sobrinus, S. criceti*, and *S. downei*), one Type IV controls competence independent of the second Type IV system (if a second system exists).

### 2.4 Integration of competence and bacteriocins varies by species

ComR boxes suggest direct regulatory targets of the ComR/XIP complex. We searched for ComR boxes in the genomes of the mutans streptococci. The predicted targets of ComR regulation fall into three categories of genes: 1.) *comS*, 2.) *comX*, and 3.) RiPP (bacteriocin) biosynthetic gene clusters (see Supplementary Table 1 for predicted ComRS regulons based on ComR boxes; see Supplementary Table 5 for analysis of the predicted RiPPs clusters using RODEO [27]). Since most mutans streptococci have two ComRS systems with only one system controlling competence, we asked which target genes are under the control of each ComRS system. To answer this question, we developed fluorescent reporters for genes of interest in each of the six transformable mutans streptococci. (See Supplementary Text 1 and Supplementary Figure 1 for details on the reporter construction.)

#### 2.4.1 S. criceti

*S. criceti* has the simplest ComRS structure of the mutans streptococci, containing a single Type IV ComRS that controls competence. Exogenous XIP increases expression of *comS*, *comX*, and the competence-associated gene *cinA* [14], [28] based on fluorescent reporters for the associated promoters (Supplementary Figure 2).

We did not find ComR boxes upstream of any bacteriocin-like gene clusters, so we hypothesize that ComRS does not directly regulate bacteriocin production in *S. criceti*. We found a *blpRH_2-like* operon with a cin-box typical of ComX regulation [14] controlling a major bacteriocin cluster in *S. criceti* as well as in *S. downei* and *S. sobrinus*, but reporters for neither the regulatory genes nor the biosynthetic genes in these clusters responded to XIP in any of the three species (data not shown).

Combined with the transformation data above, the Type IV ComRS system in *S. criceti* activates expression of *comX* and includes a positive feedback loop through *comS* activation. ComX activates competence genes (like *cinA*), but we were unable to find a bacteriocin gene cluster regulated by ComX.

#### 2.4.2 S. downei

*S. downei* has two Type IV ComRS systems, with ComRS1 and ComX required for induced competence. Reporter assays reveal that XIP1—but not XIP2—activates P_*com*S1_, P_*comX*_, and the competence gene *yfaY* (Figure 4A,B,D). XIP2 is the primary activator of P_*com*S2_ (Figure 4C). This separation of control is confirmed by the promoter responses when either ComRS system is deleted: neither XIP can induce P_*com*S1_ or P_*comX*_ when *comRS1* is deleted, and neither can induce P_*comS*2_ when *comRS2* is deleted (Supplementary Figure 3).

**Figure 4:**
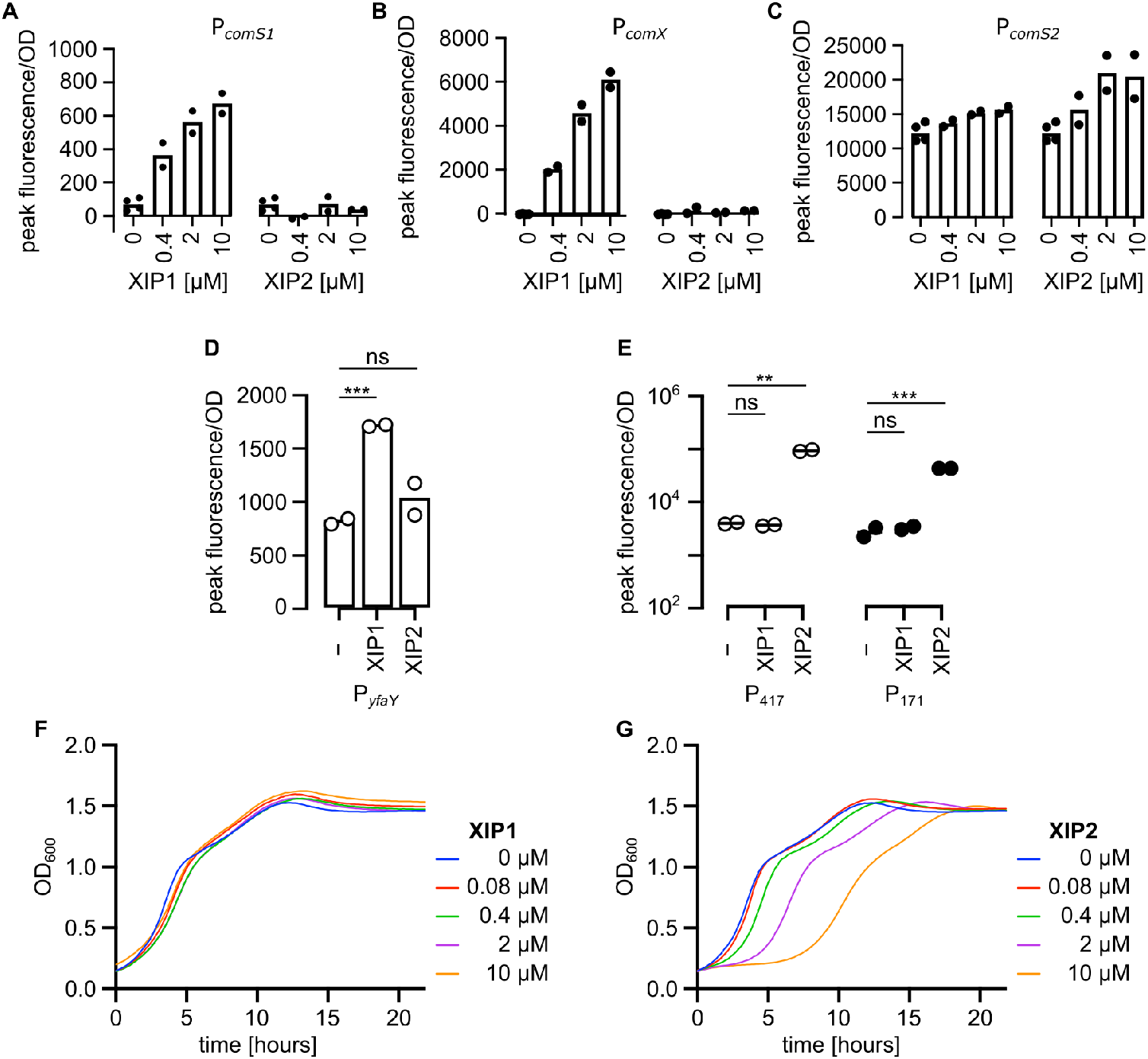
**A-B**. In *S. downei*, XIP1—but not XIP2—activates P_*comS1*_ and P_*comX*_. (By one-way ANOVA, *p* < 0.0001 for XIP1’s effect on both genes, and *p* > 0.68 and *p* > 0.13 for XIP2’s effect on P_*comS1*_ and P_*comX*_, respectively.) **C.** P_*comS2*_ is activated by both XIP1 (*p* < 0.036, ANOVA) and XIP2 (*p* < 0.032, ANOVA), but the P_*comS2*_ response is 3-fold higher with XIP2 than with XIP1 (2 *μ*M). Note that endogenous activity of P_*comS2*_ is high. **D**. XIP1 activates the competence gene *yfaY*, while **E**. XIP2 activates the bacteriocin gene clusters under the control of promoters P_417_ and P171. **F**. XIP1, which activates only competence, does not slow the growth of *S. downei* cultures. The mid-growth slowdown coincides with high P_*comS2*_, P_171_, and P_417_ (data not shown). **G**. The bacteriocin-inducing XIP2 leads to growth defect, suggesting that growth defects caused by ComRS activation in other species may be due to bacteriocins, not physiological shifts accompanying competence. The “0” and “-” conditions are DMSO controls. Significance was assessed by *t*-test (two-sided, equal variance): “ns” = *p* > 0.05; * = *p* < 0.05; ** = *p* < 0.01; *** = *p* < 0.001; **** = *p* < 0.0001.

ComR boxes are found in front of two large RiPPs biosynthetic clusters, one upstream of gene 417 (NCTC11391_00417), the first gene of a RiPPs cluster between the two *comRS* regions, and another upstream of gene 171 (NCTC11391_00171) preceding a second RiPPs cluster. We built reporter strains for the promoters of the first genes (P_417_ and P_171_). XIP2 activated both of these promoters, but XIP1 did not (Figure 4E). It appears that *S. downei* independently controls competence and bacteriocin production using ComRS1 and ComRS2, respectively.

Adding exogenous XIP temporarily slows growth in many streptococci. The growth arrest may be caused by a redirection of cellular machinery toward competence or because of induced self-acting bacteriocins (as observed in *S. mutans, S. thermophilus*, and *S. sanguinus* [15], [16], [23]). The independence of the competence-controlling ComRS1 and the bacteriocin-controlling ComRS2 in *S. downei* allowed us to test the relative effects of each process on growth. XIP1 has no effect on *S. downei*’s growth, while XIP2 creates a dosedependent growth defect (Figure 4F,G). In fact, the inherent mid-growth slowdown before the stationary phase in *S. downei—*even without exogenous XIP—coincides with the high activity phases of P_*comS*2_, P_171_, and P_417_. Thus in *S. downei*, bacteriocins—not competence—underlie the XIP-induced growth defect.

#### 2.4.3 S. sobrinus

We previously discovered two *comR* genes in *S. sobrinus, comR1* and *comR2* [21]. Unlike *S. downei*, only *comR1* in *S. sobrinus* has an associated *comS*, while *comR2* does not (Figure 5A). However, ComR2 is required for competence, and ComR1 inhibits transformation efficiency by an unknown mechanism [21]. We also found that *S. sobrinus*’ XIP activates bacteriocins that inhibit the growth of *S. mutans*—an effect that disappeared when the gene cluster containing *comR1, comS* and *comR2* was deleted [21].

**Figure 5:**
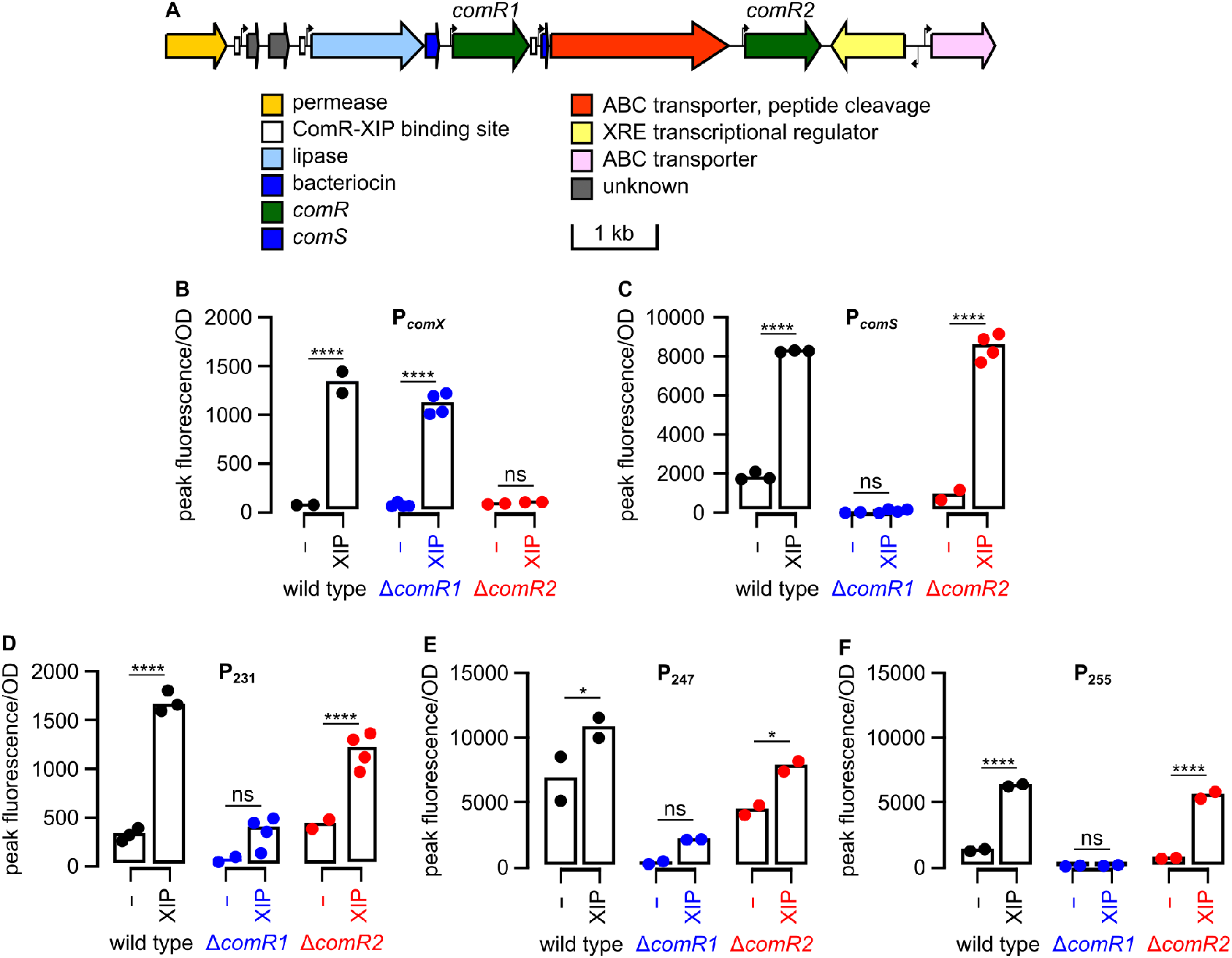
**A**. *S. sobrinus* has two *comR* genes in the same gene cluster. The only *comS* is immediately downstream of *comR1*. **B**. ComR2— but not ComR1—is required for *comX* activation. We previously showed that ComR2 is also necessary for competence [21]. **C**. Only ComR1 creates a feedback loop through ComS, suggesting the ComR box preceeding *comS* is bound only by ComR1/XIP. Three ComR box-containing promoters in the bacteriocin cluster upstream of *comR1* (P231 in panel **D**, P247 in panel **E** and P255 in panel **F**) are controlled by ComR1. The *comR1* deletion abolishes their endogenous activities in addition to their response to XIP. Significance was assessed by *t*-test (two-sided, equal variance): “ns” = *p* > 0.05; * = *p* < 0.05; **** = *p* < 0.0001.

Since *S. sobrinus* has only one ComS/XIP, we used gene deletion strains and a P_*comX*_ reporter to identify the ComR that activates *comX*. XIP induced P_*comX*_ expression in a *ΔcomR1* strain but not in a *ΔcomR2* strain (Figure 5B), even at elevated XIP concentrations (Supplementary Figure 3A). As expected, only the competence-controlling ComR2 activates ComX.

In *S. downei*, both ComR proteins activate their associated P*comS* to create positive feedback loops. The only *comS* gene in *S. sobrinus* is downstream of *comR1* (Figure 5A), and reporter assays show that only ComR1 appreciably activates P_*comS*_ even though ComR1 was shown to inhibit competence (Figure 5C) [21]. In a *ΔcomR1* background, XIP creates a weak dose-dependent response of P_*comS*_ through ComR2 (Supplementary Figure 3B); however, the maximum activity of P_*comS*_ in a *ΔcomR2* strain is 39-fold higher than in a *ΔcomR1* strain (Figure 5C). Taken together, it appears that the endogenous positive feedback loop of XIP expression is achieved through ComR1, even though ComR2 is the protein that induces competence in response to the same XIP [21].

A bacteriocin cluster similar to the *S. downei* 417 cluster sits upstream of *S. sobrinus comR1* [21]. To determine if this cluster is controlled by ComR1 or ComR2, we constructed fluorescent reporters for the three ComR-box containing promoters in this bacteriocin cluster (P231, P247, and P255). All three reporters are induced by exogenous XIP, but only in strains with *comR1* (Figure 5D,E,F). Deleting *comR2* had no significant effect on the activity of these promoters. Thus ComR1 regulates the 417-like bacteriocin cluster.

In summary, *S. sobrinus* has two Type IV ComR systems that regulate independent sets of genes despite being activated by the same XIP. ComR1 is forms a positive feedback loop for ComS/XIP and activates a bacteriocin gene cluster, while ComR2 activates ComX to control competence.

Homology suggests that both ComR1 and ComR2 in *S. sobrinus* are more related to the bacteriocin-regulating *S. downei* ComR2 than the competence-controlling *S. downei* ComR1 (Figure 2). This is confirmed by alignment of the ComRS genomic regions in *S. sobrinus* and *S. downei* (Supplementary Figure 5), which also shows that the genomic region including *comRS1* in *S. downei* is missing in *S. sobrinus*. We previously hypothesized that a duplication event to restore competence in *S. sobrinus* would explain why the two ComR proteins in *S. sobrinus* share a single XIP ([21], Supplementary Figure 5). The lone ComRS in *S. criceti*, which controls competence, is more similar to the competence-controlling ComRS1 in *S. downei* (Supplementary Figure 5); in fact, both ComR proteins share the same XIP sequence (Table 1). By contrast, the region directly downstream of *S. downei comRS1* contains both a bacteriocin cluster and a bacteriocin-controlling *comRS2*, but this region is missing in *S. criceti* (Supplementary Figure 5). Our previous analysis of the ComR regions of three *S. sobrinus* strains also revealed three different genomic structures [21]. These alignment results indicate high frequency genomic recombination in the ComR region, likely facilitated by two nearby *comR* genes. This remodeling changes the competence-bacteriocin network topology.

#### 2.4.4 S. mutans

The Type IV ComR2 was originally described as an Rgg-type transcriptional regulator and was hypothesized to activate a nearby bacteriocin gene cluster (SMU.370–SMU.379) [11]. Indeed, the XIP2 derived from the ComS2 we discovered near ComR2 activates a reporter for SMU.370, the first gene in the bacteriocin cluster (Figure 6A), in addition to activating *comS2*. XIP2 does not increase the peak activities of the promoters for *comX*, the competence gene *cinA*, the bacteriocin regulator *blpR*, or bacteriocin gene *nlmA* (Figure 6B). Most of these reporters were induced by XIP1 (Figure 6B), corroborating previous studies [13], [14].

**Figure 6:**
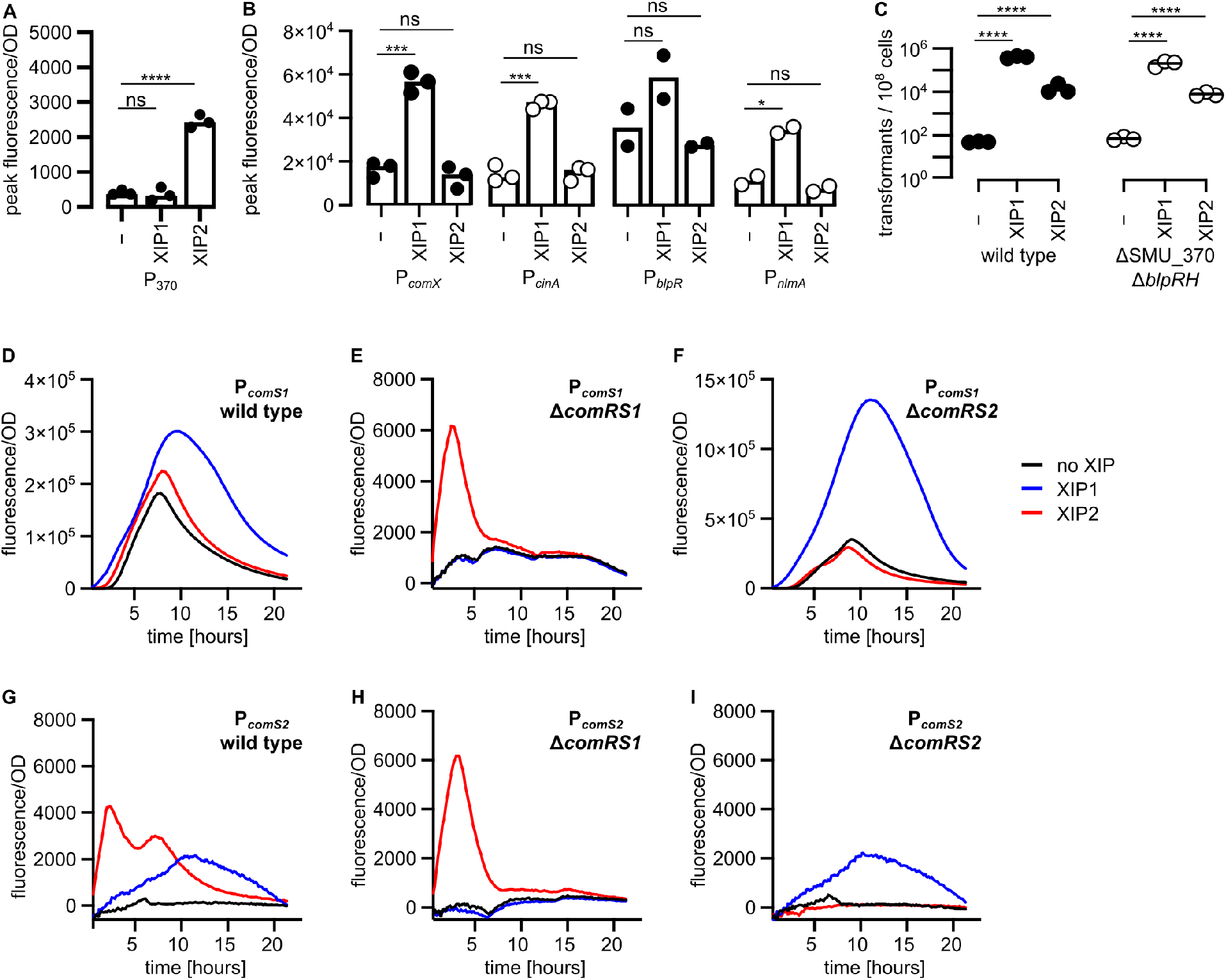
**A**. The newfound Type IV ComRS regulates a promoter upstream of the nearby bacteriocin gene cluster, as originally hypothesized in [11] before the discovery of the Type IV ComS. **B**. XIP2 in *S. mutans* does not increase the peak activity of *comX*, the competence gene *cinA*, or the bacteriocin regulator *blpR* and bacteriocin gene *nlmA*. Most of these reporters were activated when XIP1 was added to the media (*blpR* was reported to be induced by XIP1 [13], [14]; the lack of significance here is likely due to low replicate number). **C**. Although the Type IV ComRS2 does not activate *comX* or competence genes, XIP2 induces synthetic competence at ~100× lower levels than XIP1. Deleting bacteriocins controlled by ComR2 (SMU_370–SMU_379) and the ComDE system controlled by ComR1 does not affect the XIP2-induced competence, suggesting the competence may be due to crosstalk between ComR1 and ComR2, instead of bacteriocin effects on cell permeability and ComS/XIP release. Bars in panels **A** and **B** and lines in panel **C** are the averages of three biological replicates. **D**. XIP1 creates a broad peak of ComS1 expression centered after ~12 hours. **E.** Removing the ComRS1 pathway reveals a XIP2-induced smaller, narrow peak at ~4 hours. Note the difference in scale between panels **D** and **E**. **F**. Deleting ComRS2 does not affect the large, broad peak in ComRS1. **G**. Upon XIP2 addition, P_*comS2*_ activity shows two peaks—an early peak caused by XIP2 through ComRS2 (**H**) and a smaller, broader peak caused by crosstalk from XIP1 (**I**). Times in panels **D**–**I** measure the time since the addition of XIP1, XIP2, or a DMSO control. Traces are the average of two biological replicates. Significance was assessed using log_10_-transformed values by *t*-test (two-sided, equal variance): “ns” = p > 0.05; * = *p* < 0.05; *** = *p* < 0.001; **** = *p* < 0.0001.

At first glance, the Type II and Type IV ComRS systems in *S. mutans* appear independent. The Type II ComRS1 activates ComX, competence, ComCDE, and subsequently bacteriocins. The Type IV ComRS2 activates a separate cluster of bacteriocin genes [14]. However, we observed crosstalk between ComRS1 and ComRS2 in *S. mutans*. Our first evidence was the transformation data where XIP2 induced competence but required both ComRS1 and ComRS2 (Figure 3E).

We tested two hypotheses for how ComRS2 can activate competence without directly regulating ComX: bacteriocin-enhanced XIP1 transport, or crosstalk between ComRS1 and ComRS2. Others have suggested that bacteriocins can potentiate competence by creating pores in the cell membrane that increase the diffusion of XIP into the cell [13], [23], or by lysing cells to release intracellular ComS/XIP [29]. We deleted the bacteriocin cluster activated by ComR2, but neither this deletion nor the subsequent deletion of *blpCRH* reduced competence from either XIP1 or XIP2 (Figure 6C). It is possible that ComRS2 regulates other bacteriocins in *S. mutans*, but a bioinformatic search for ComR2 boxes did not reveal any additional candidate gene clusters.

Our second hypothesis is that ComRS2 increases the expression of *comS1*, thereby initiating the ComRS1 positive feedback loop that drives activation of *comX*. A P_*comS*1_ reporter strain shows that XIP2 does not significantly increase *peak P*_*comS*1_ activity above background levels, possibly due to the high endogenous ComRS1 activity (Figure 6D, Supplementary Figure 6). In a *ΔcomRS1* background, however, XIP2 induces P_*comS*1_ activity 4-fold above background (*p* = 0.003, *t*-test). This activation requires ComRS2 (Figure 6E–F). ComRS2’s activation of P_*com*S1_ is comparatively small, but it occurs faster than the on-pathway peak in ComS1 expression. In a *ΔcomRS1* strain, P_*comS*1_ activity peaks within 3 hours of XIP2 addition (Figure 6E). The larger P_*comS*1_ peak in a wild type background occurs 7–9 hours after XIP1 addition (Figure 6D). It appears that ComRS2 can “kickstart” the ComRS1 feedback loop by activating *comS1* expression early on. This kickstarting effect becomes visible in wild-type *S. mutans* in the early phase of P_*comS*1_ activity, especially when XIP2 is added at low OD before endogenous ComRS1 activity peaks (Supplementary Figure 6).

Conversely, we found that ComRS1 can activate the ComRS2 system. Addition of XIP1 leads to a delayed, broad peak of P_*comS*2_ activity that coincides with high P_*comS*1_ activity. XIP1’s effect on P_*com*S2_ requires ComRS1 (Figure 6G,H). The cross-activation of P_*comS*2_ by XIP1 is comparable to on-pathway activation levels by ComR2/XIP2 (Figure 6I,G). This creates bimodal P_*comS*2_ activity when XIP2 is added: a fast, larger peak through ComRS2, and a second broader peak caused by crosstalk from ComRS1 (Figure 6G).

In summary, crosstalk exists in both directions between ComRS1 and ComRS2 in *S. mutans*. Competence and the BlpCRH bacteriocin system downstream of ComX are primarily controlled by ComRS1, and ComRS2 controls a separate bacteriocin cluster. ComRS1 activity sills over and activates the ComRS2 system, while the ComRS2 system can kickstart the positive feedback loop of the ComRS1 system.

#### 2.4.5 *S. ratti* and *S. ferus*

*S. ratti*, like *S. mutans*, has both Type II and Type IV ComRS systems. The Type II ComRS1 activates ComX, but the Type IV ComRS2 does not (Supplementary Figure 7A). The same is true in *S. ferus;* however, *S. ferus* was the only species where adding XIP2 did not lead to activation of a P_*comS*2_ reporter (Supplementary Figure 7B). Two short peptide-encoding genes appear downstream of the *S. ferus comR2* gene: *comS2a* and *comS2b*. Neither of the predicted XIP peptides activated the predicted P_*comS*2_ (Supplementary Figure 7B). We are unsure if either of these ORFs encode the true ComS2.

*S. ratti* also shows crosstalk between ComRS1 and ComRS2 similar to *S. mutans*. Such crosstalk appears in our reporter data (Supplementary Figure 8) but not in our transformation data. XIP2 in *S. ratti* induces a fast, sharp peak of P_*comS*1_ activity and a subsequent larger peak; the former is likely caused by ComRS2, and the latter is from ComRS1 due to its positive feedback loop being kickstarted by ComRS2. XIP1 cross-induces P_*comS*2_ with a broader, later peak, although the induction is tenfold lower than on-pathway induction by XIP2. This wiring differs from the crosstalk in *S. mutans* (Supplementary Figure 8).

### 2.5 ComR recognition sequences confer specificity of Type II and Type IV ComRS systems

We were intrigued by how paralogous ComRS systems insulate their signals within the same cell. Previous work showed how small changes in the ComR box sequence can ablate ComR recognition [15], [17]. We tested if we can switch the recognition of P_*comX*_ by ComR1 and P_*comS*2_ by ComR2 by creating promoter variants with altered ComR boxes. (We avoided using P_*comS*1_ since its higher strength may mask crosstalk in wild-type background.) P_*comX*_ and P_*comS*2_ have three differences among their conserved nucleotides [15] and four differences at other positions in their 20 bp ComR boxes (Figure 7A, see Supplementary Figure 9A for full analysis). In wild type strains, XIP1 induces a high, broad peak of P_*comX*_ activity at 12 hours (Figure 7B) while XIP2 induces a lower, sharper peak of P_*comS*2_ activity at ~2 hours (Figure 7D).

**Figure 7:**
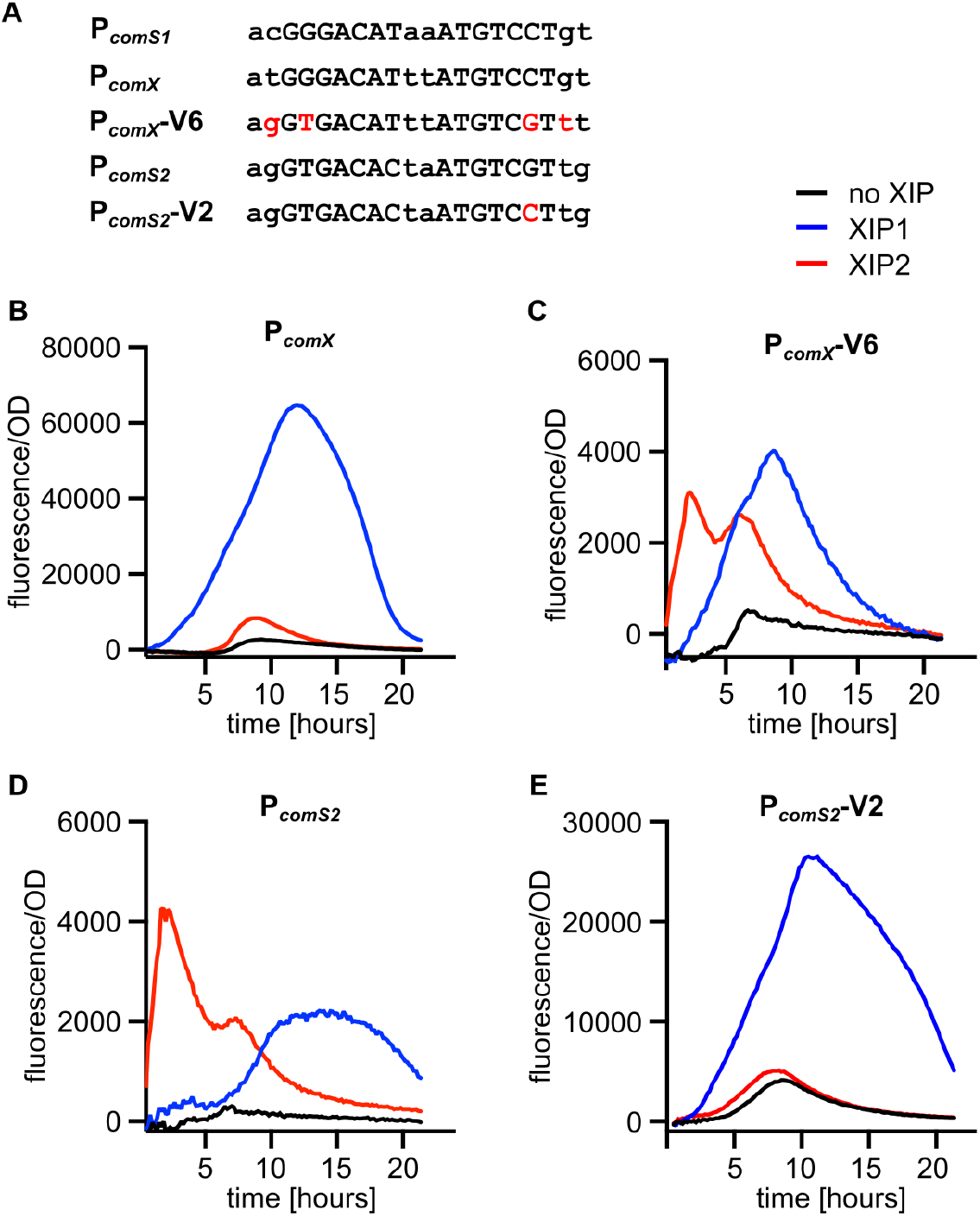
A small number of base pairs in ComR boxes confer specificity for the ComRS1 or ComRS2 pathways in *S. mutans*. **A**. Promoters for *comX* and *comS2* are targeted by ComRS1 and ComRS2, respectively. We constructed variants of both promoters by changing nucleotides in P_*comS2*_ to match those in P_*comX*_, and vice versa. Capitalized bases indicate the most conserved positions of ComR boxes [15]. **B**. XIP1 induces a broad, delayed peak of P_*comX*_ activity; effect by XIP2 is largely masked by endogenous activity. **C**. The P_*comX*_-V2 variant shows a sharp, early peak from XIP2 and a smaller peak from XIP1 (note the difference in scale between **B** and **C**). **D**. The sharp XIP2 peak and small XIP1 response typify the P_*comS2*_ response, suggesting that the V2 variant of P_*comX*_ behaves similar to P_*comS2*_. **E**. Changing position 17 of P_*comS2*_ to match P_*comX*_ makes the P_*comS2*_-V2 response similar to the P_*comX*_ promoter—a broad, late peak induced by XIP1, and little induction by XIP2. Traces are the averages of two biological replicates.

A single `G`_17_ → `C` mutation in P_*comS*2_ (P_*comS*2_-V2) switched its response profile from XIP2-type to XIP1-type. The promoter variant lost the XIP2-induced sharp peak and gained the XIP1-induced large peak (Figure 7E). In contrast, changing four bases in P_*comX*_ (P_*comX*_-V6) makes the promoter’s response profile similar to that of P_*comS2*_—a sharp, early peak induced by XIP2 and an attenuated, later peak from XIP1 (Figure 7C, Supplementary Figure 9C–I). A T_4_ → `G` substitution in P_*comS*2_ makes it unresponsive to either XIP, while changing `G`_4_ → `T` in P_*comX*_ did not affect its response profile (Supplementary Figure 9). Taken together, the specificity of the *S. mutans* Type II and Type IV ComRS systems can be switched by changing the ComR boxes upstream of the target genes. Making the ComRS2 targets responsive to ComRS1 would require a single mutation, while making the ComRS1 targets responsive to ComRS2 requires four mutations.

We also tried to switch the regulation of P_*comS*_ or P_*comX*_ in *S. sobrinus* from ComR1 to ComR2 and vice versa. Mutations either do not change the ComR1/ComR2 specificity or make the reporter gene unresponsive to both regulators (Supplementary Figure 10). Swapping the ComR boxes upstream of the *comX* and *comS* genes also creates unresponsive reporters (Supplementary Figure 10). It appears that specificity in *S. sobrinus* is not entirely controlled by the ComR boxes, and the targets of the ComR1 and ComR2 are better insulated from crosstalk.

## 3 Discussion

New ComRS systems practically and conceptually change our understanding of the mutans streptococci. Practically, the competence-controlling ComRS systems allow genetic manipulation of the previously intractable *S. ratti*, *S. ferus*, *S. downei*, and *S. criceti*; *S. macacae* remains the only mutans streptococcus that cannot be transformed in our hands despite having both a Type II and Type IV ComRS. Natural competence opens the door for comparative studies of this important group of bacteria. Our work highlights the value of such comparative analysis by showcasing the diverse functions of the ComRS system. The Type IV ComRS systems that control bacteriocins are new tools for activating these natural products and studying antimicrobial defenses among the oral microbes.

Conceptually, our discovery of dual ComRS systems changes our view of this important pathway. With the exception of *S. criceti*, all mutans streptococci have two ComRS systems: competence is controlled primarily by one system (together with bacteriocins in *S. mutans*), while the other system primarily controls bacteriocin synthesis (Figure 8). Species in the mutans subclade use a Type II ComRS to regulate competence and a Type IV ComRS to regulate bacteriocins, while species in the sobrinus subclade use two Type IV systems to regulate either competence or bacteriocins (Figure 8; see Supplementary Figure 11 for more detailed network structures). Such separation may not be unique to the mutans group. In *S. pneumoniae*, the BlpRH pathway can activate bacteriocins independent of competence-controlling ComCDE pathway [18]–[20]. Mignolet, *et al.* recently identified ComR paralogs in *S. salivarius* that may be dedicated to controlling bacteriocins, although they could not find a corresponding peptide pheromone encoded in the genome [17].

**Figure 8:**
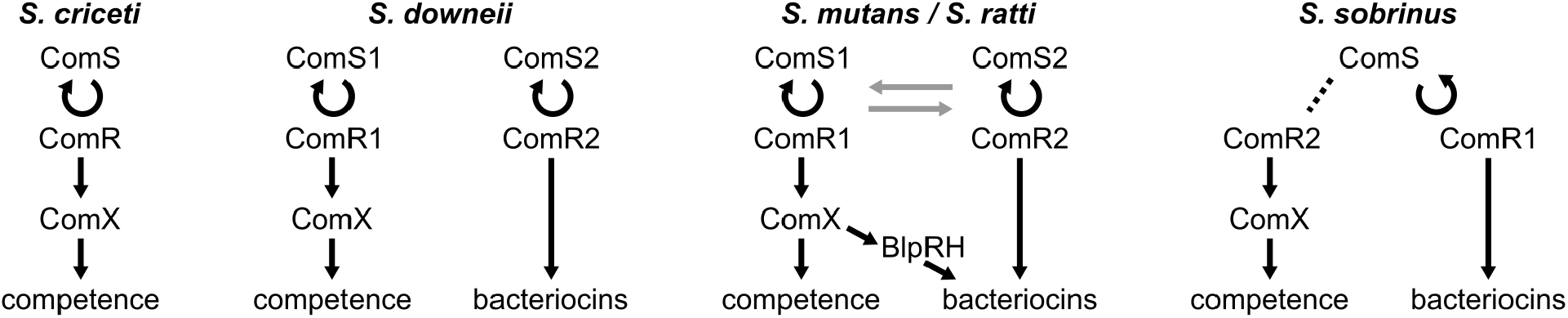
ComRS pathways in the oral streptococci show differing levels of crosstalk in the regulation of competence and bacteriocins. Gray lines for *S. mutans* and *S. ratti* indicate crosstalk between the Type II and Type IV pathways. Note that ComX’s control of bacteriocins applies to *S. mutans* but not *S. ratti*, as neither *blpRH* homologs nor ComX control of bacteriocins are found in *S. ratti*. The dotted line in *S. sobrinus* indicates that ComR2 binds with ComS/XIP but does not activate *comS* to form a positive feedback loop.

However, *S. sobrinus* and *S. mutans*, for example, do not cleanly separate their ComRS systems. *S. sobrinus* uses a single XIP to activate both ComR proteins, effectively integrating the two systems (Figure 8). The ComRS1 system in *S. mutans* can activate the ComRS2 system during its high activity phase, while the ComRS2 system can kickstart ComRS1 and induce low levels of competence. This is reminiscent of the bidirectional crosstalk in the ComCDE-BlpCRH network of *S. pneumoniae* [18]–[20]. The similarity in topology is striking, especially considering the different molecular mechanisms used by these two species. Unlike the ComRS pathway, ComCDE and BlpCRH in *S. pneumoniae* are two-component systems, and BlpCRH stimulates ComCDE through transporter promiscuity [18]–[20]. In both cases, one system controls both competence and bacteriocins, and another paralogous system primarily controls only bacteriocins, while the two systems are neither fully orthogonal nor fully integrated, but have intermediate levels of crosstalk. Such convergence of network topology leads to questions about its potential advantages for the bacteria.

Multiple ComRS pathways with crosstalk creates an array of network structures and dynamics in the mutans group streptococci (Figure 8). The network diversity suggests evolutionary plasticity whereby networks can be re-wired based on environmental conditions. Such plasticity is made possible by maintaining paralogous regulatory systems, since similarity in both the signaling proteins and their binding sites increases the probability of rewiring. In *S. mutans*, changing a single base pair in the ComR box can switch the network topology for a set of target genes (Figure 7). Similar cases have been observed in *S. pneumoniae* and *S. salivarius* [17], [20].

More broadly, the streptococcal competence/bacteriocin networks exhibit modularity, weak regulatory linkage, and reduced pleiotrophy—all hallmarks of facilitated variation [30]. Other researchers have proposed facultative cheating as mechanism for maintaining paralogous quorum-sensing systems [31], [32]. Variable and stressful environmental conditions like those found in the oral microenvironment could also favor the genome plasticity we see in the mutans streptococci.

This study used fluorescent reporter strains to study ComRS systems *in vitro*. The oral streptococci live in the supragingival plaques of mammalian teeth and experience a wider array of nutrients and stressors than our chemically defined media. The oral microenvironment and physical contact with other cells in the biofilm can alter the behavior of the ComRS systems [7]. Our assays measured the average responses to populations of cells, but ComRS systems can behave stochastically on the single-cell level [29]. A systems-level analysis that combines environmental and social factors and stochastic effects is necessary to fully understand the intra- and interspecies dynamics of the ComRS pathways.

## 4 Methods

### 4.1 Strains, reagents, and culture conditions

Full lists of primers, synthesized oligonucleotide blocks, plasmids, and strains appear in Supplementary Tables 2, 3, and 4. Type strains were purchased from ATCC (*S. mutans* UA159 and *S. sobrinus* SL1) or DSMZ (*S. ratti* FA1, *S. ferus* 8S1, *S. macacae* 25-1, *S. downei* MFe28/NCTC 11391, *S. criceti* HS-6, and *S. sobrinus* NCTC 10919). *S. sobrinus* NIDR 6715-7 was provided by the ARS Culture Collection (NRRL B-14554).

Cells were grown in either Todd Hewlitt Broth + 0.5% yeast extract (Sigma), a chemically defined medium (CDM) [33], [34], or a mixture of both as described below. Solid media were made by adding 1.5% agar. All cultures were grown at 37 °C. *S. sobrinus* was transformed anaerobically (5% H_2_, 10% CO_2_, and 85% N_2_) in an anaerobic chamber (Coy Labs, MI, USA). All other species were grown and transformed aerobically under 5% CO_2_. Antibiotics (Sigma) were used with selectable markers: 1 mg/ml kanamycin, 400 *μ*g/ml spectinomycin, or 4 *μ*g/ml chloramphenicol for streptococci; 100 *μ*g/ml spectinomycin for *Escherichia coli*.

Enzymes were purchased from New England Biolabs (Ipswich, MA, USA). Oligonucleotides were synthesized by Integrated DNA Technologies (Coralville, IA, USA). Synthetic peptides were purchased from GenScript, Inc. (Piscataway, NJ, USA) at >90% purity.

### 4.2 Transformation assays

For strain construction in all species except *S. sobrinus*, overnight cultures were grown in CDM (or 1:1 THY+CDM for *S. mutans* to improve growth). The next day, cultures were diluted 150× into pre-warmed CDM and grown to OD 0.5–0.8 (Biochrom CO8000). Cultures were vortexed with the corresponding XIP (2 *μ*M final concentration) before moving 50 *μ*l of culture into a 0.5 ml centrifuge tube and vortexing with DNA (100 ng of plasmid or 5 *μ*l of Golden Gate reaction mix). Cells were incubated for 2 hours at 37 °C under 5% CO_2_. Afterwards, 3–10 *μ*l of the transformation mix was spread by glass beads on THY agar plates with the appropriate antibiotic. For *S. sobrinus* strains—which have a lower transformation efficiency—cultures were grown anaerobically, 10 *μ*M XIP was added to 200 *μ*l culture, and the entire 200 *μ*l was plated for selection. Colonies were picked after 18–30 hours of incubation.

To measure transformation efficiency, overnight cultures were grown in THY to suppress endogenous competence induction. The next day, cultures were diluted 150× into pre-warmed CDM, and grown to OD 0.5-0.8. Using a multichannel pipette, 100 *μ*l of culture was then added to each well of a 96 well plate. XIPs diluted in DMSO or an equal volume of DMSO was added to a final concentration of 2 *μ*M, along with 100 ng of plasmid pRW49 in water. The plasmid pRW49 contains the *aad9* spectinomycin resistance gene and a gene encoding the fluorescence protein mScarlet-1. The plate was shaken for 5–10 s with a plate mixer (Genie SI-0400A Digital MicroPlate), taking care to avoid splashing between wells. After two hours of incubation (37 °C, 5% CO_2_), the transformation mixes were diluted serially from 10^-1^ to 10^-6^, and 10 ul from each well was spotted onto 10×10 cm square agar plates with or without spectinomycin. Colonies were counted after 24–30 hours.

### 4.3 Plasmid construction

Generic reporters were constructed using Golden Gate assembly [35] and plasmid pRW17 [21]. Synthesized gBlocks containing promoter sequences were combined with fluorescent reporter genes codon-optimized for *S. mutans*. Promoters were at least 150 bp immediately upstream of the start codon and included the native RBS for each gene. The constructed reporters are seamless, having no extra bases inserted between the RBS and start codon of the reporter genes. The expression cassette is insulated from the rest of the plasmid by two bidirectional terminators. The Golden Gate reaction was used to transform *E. coli* DH5*α*. Plasmids were verified by Sanger sequencing before transformation into the streptococci.

### 4.4 Gene deletions

Fragments (350–100 bp) flanking the gene to be deleted were PCR amplified and ligated to an antibiotic resistance marker by Golden Gate assembly. Transformants were verified by colony PCR and Sanger sequencing.

### 4.5 Fluorescent reporter assays

Overnight cultures were grown in a 1:1 mixture of CDM and THY in5%CO_2_ incubator. (The THY suppresses ComRS activation.) The next day, the cultures were diluted 250× into 250 ul prewarmed CDM in a black-walled 96-well plate and grown for ~3 hours (OD between 0.1 and 0.15 including 0.85 of background OD, corresponding to around 0.1 of normal OD) in a Cytation 5 plate reader with 5% CO_2_ at 37 °C. Growth and fluorescence were measured every 10 minutes following 10 s of shaking. The excitation/emission wavelengths (nm) used for each fluorescent protein are: 500/520 for mNeonGreen and sfGFP; 569/593 for mScarlet-I; 462/492 for mTFP1; 399/454 for mTagBFP2; and 434/474 for mTurquoise2. After the 3 hours of incubation 1.25 *μl* of 0.4 mM XIP or an equal volume of DMSO were quickly added to a final concentration of 2 *μ*M, unless otherwise specified. At least two biological replicates were performed for each condition. Incubation and readings continued for at least 20 hours.

### 4.6 Reporter data processing

Background fluorescence from media decreases nonlinearly as cell culture OD increases [36]. Strains without a fluorescent reporter protein were included for all experiments. Readings from these strains were fitted to a second-order liner model using OD, the time derivative of OD, and t as features. Models were fit using the PolynomialFeatures transformer and LinearRegression estimator in scikit-learn Python package [37]. The predicted background fluorescence was subtracted from experimental wells before processing.

Fluorescence/OD (FL/OD) readings were filtered with a Savitzky-Golay filter to smooth noise (window length of 41; 4th-order for *S. mutans* and 3rd order otherwise). The filter parameters were selected manually to avoid overfitting the initial (low OD) values and avoid underfitting low accuracy peaks. An affine shift of 0.114 OD was added to dampen noise in FL/OD at early timepoints with low OD. The peak FL/OD was found as a local maximum after the initial region of decreasing FL/OD was removed to prevent maxima caused by noise at early timepoints. The FL/OD was averaged at five adjacent points to determine the peak fluorescence. The activation time of a peak is defined as the timepoint when the derivative of FL/OD reached 25% of its maximal value.

## 5 Acknowledgements

We thank Bill Metcalf, Shannon Sirk, and Pablo Perez-Pinera for their feedback and suggestions; Jiayi Cai for help in reporter data processing with Python; Aynur Namik for assistance in experiments. This work was supported by the National Institutes of Health grant GM138210.

## Notes

### Competing Interest Statement

The authors have declared no competing interest.

## References

[1] M. Blokesch, “Natural competence for transformation,” Current Biology, vol. 26, no. 21, R1126–R1130, Nov. 2016, issn: 0960-9822. doi: 10.1016/j.cub.2016.08.058.

[2] C. Johnston, B. Martin, G. Fichant, P. Polard, and J.-P. Claverys, “Bacterial transformation: Distribution, shared mechanisms and divergent control,” Nature Reviews Microbiology, vol. 12, no. 3, pp. 181–196, Mar. 2014, issn: 1740-1534. doi: 10.1038/nrmicro3199.

[3] J. C. Mell and R. J. Redfield, “Natural Competence and the Evolution of DNA Uptake Specificity,” Journal of Bacteriology, vol. 196, no. 8, pp. 1471–1483, Apr. 2014, issn: 0021-9193. doi: 10.1128/JB.01293-13.

[4] E. Shanker and M. J. Federle, “Quorum Sensing Regulation of Competence and Bacteriocins in Streptococcus Pneumoniae and Mutans,” Genes, vol. 8, no. 1, Jan. 2017, issn: 2073-4425. doi: 10.3390/genes8010015.

[5] L. Fontaine, A. Wahl, M. Fléchard, J. Mignolet, and P. Hols, “Regulation of competence for natural transformation in streptococci,” Infection, Genetics and Evolution, vol. 33, pp. 343–360, Jul. 2015, issn: 1567-1348. doi: 10.1016/j.meegid.2014.09.010.

[6] H. Sztajer, S. P. Szafranski, J. Tomasch, et al.,“Cross-feeding and interkingdom communication in dual-species biofilms of Streptococcus mutans and Candida albicans,” The ISME Journal, vol. 8, no. 11, pp. 2256–2271, Nov. 2014, issn: 1751-7370. doi: 10.1038/ismej.2014.73.

[7] J. R. Kaspar, K. Lee, B. Richard, A. R. Walker, and R. A. Burne, “Direct interactions with commensal streptococci modify intercellular communication behaviors of Streptococcus mutans,” The ISME Journal, vol. 15, no. 2, pp. 473–488, Feb. 2021, issn: 1751-7370. doi: 10.1038/s41396-020-00789-7.

[8] C. A. Werlang, W. G. Chen, K. Aoki, et al.,“Mucin O-glycans suppress quorum-sensing pathways and genetic transformation in Streptococcus mutans,” Nature Microbiology, vol. 6, no. 5, pp. 574–583, May 2021, issn: 2058-5276. doi: 10.1038/s41564-021-00876-1.

[9] Y.-H. Li, X.-L. Tian, G. Layton, C. Norgaard, and G. Sisson, “Additive attenuation of virulence and cariogenic potential of Streptococcus mutans by simultaneous inactivation of the ComCDE quorum-sensing system and HK/RR11 two-component regulatory system,” Microbiology (Reading, England), vol. 154, no. Pt 11, pp. 3256–3265, Nov. 2008, issn: 1350-0872. doi: 10.1099/mic.0.2008/019455-0.

[10] L. Fontaine, C. Boutry, M. H. de Frahan, et al.,“A Novel Pheromone Quorum-Sensing System Controls the Development of Natural Competence in Streptococcus Thermophilus and Streptococcus Salivarius,” Journal of Bacteriology, vol. 192, no. 5, pp. 1444–1454, Mar. 2010, issn: 1098-5530. doi: 10.1128/JB.01251-09.

[11] L. Mashburn-Warren, D. A. Morrison, and M. J. Federle, “A Novel Double-Tryptophan Peptide Pheromone Controls Competence in Streptococcus Spp. via an Rgg Regulator.,” Molecular Microbiology, vol. 78, no. 3, pp. 589–606, Nov. 2010. doi: 10.1111/j.1365-2958.2010.07361.x.

[12] D. A. Morrison, R. Khan, R. Junges, H. A. Amdal, and F. C. Petersen, “Genome Editing by Natural Genetic Transformation in Streptococcus Mutans.,” Journal of microbiological methods, vol. 119, pp. 134–141, Dec. 2015. doi: 10.1016/j.mimet.2015.09.023.

[13] M. Reck, J. Tomasch, and I. Wagner-Döbler, “The Alternative Sigma Factor SigX Controls Bacteriocin Synthesis and Competence, the Two Quorum Sensing Regulated Traits in Streptococcus mutans,” PLOS Genetics, vol. 11, no. 7, e1005353, Jul. 2015,issn: 1553-7404. doi: 10.1371/journal.pgen.1005353.

[14] R. Khan, H. V. Rukke, H. Hovik, et al.,“Comprehensive Transcriptome Profiles of Streptococcus mutansUA159 Map Core Streptococcal Competence Genes,” mSystems, vol. 1, no. 2, M. J. McFall-Ngai, Ed., e00038–15, Apr. 2016. doi: 10.1128/mSystems.00038-15.

[15] L. Fontaine, P. Goffin, H. Dubout, et al.,“Mechanism of competence activation by the ComRS signalling system in streptococci,” Molecular Microbiology, vol. 87, no. 6, pp. 1113–1132, 2013, issn: 1365-2958. doi: 10.1111/mmi.12157.

[16] J. Mignolet, L. Fontaine, A. Sass, et al.,“Circuitry Rewiring Directly Couples Competence to Predation in the Gut Dweller Streptococcus Salivarius,” Cell Reports, vol. 22, no. 7, pp. 1627–1638, Feb. 2018, issn: 2211-1247. doi: 10.1016/j.celrep.2018.01.055.

[17] J. Mignolet, G. Cerckel, J. Damoczi, et al.,“Subtle selectivity in a pheromone sensor triumvirate desynchronizes competence and predation in a human gut commensal,” eLife, vol. 8, 2019, issn: 2050-084X. doi: 10.7554/eLife.47139.

[18] W.-Y. Wholey, T. J. Kochan, D. N. Storck, and S. Dawid, “Coordinated Bacteriocin Expression and Competence in Streptococcus pneumoniae Contributes to Genetic Adaptation through Neighbor Predation,” PLOS Pathogens, vol. 12, no. 2, e1005413, Feb. 2016, issn: 1553-7374. doi: 10.1371/journal.ppat.1005413.

[19] M. Kjos, E. Miller, J. Slager, et al.,“Expression of Streptococcus pneumoniae Bacteriocins Is Induced by Antibiotics via Regulatory Interplay with the Competence System,” PLOS Pathogens, vol. 12, no. 2, e1005422, Feb. 2016, issn: 1553-7374. doi: 10.1371/journal.ppat.1005422.

[20] C. Y. Wang, N. Patel, W.-Y. Wholey, and S. Dawid, “ABC transporter content diversity in Streptococcus pneumoniae impacts competence regulation and bacteriocin production,” Proceedings of the National Academy of Sciences of the United States of America, vol. 115, no. 25, E5776–E5785, Jun. 2018, issn: 1091-6490. doi: 10.1073/pnas.1804668115.

[21] J. W. Li, R. M. Wyllie, and P. A. Jensen, “A Novel Competence Pathway in the Oral Pathogen Streptococcus sobrinus,” Journal of Dental Research, vol. 100, no. 5, pp. 542–548, May 2021, issn: 1544-0591. doi: 10.1177/0022034520979150.

[22] H. Steinmoen, A. Teigen, and L. S. Håvarstein, “Competence-Induced Cells of Streptococcuspneumoniae Lyse Competence-Deficient Cells of the SameStrain duringCocultivation,” Journal of Bacteriology, vol. 185, no. 24, pp. 7176–7183, Dec. 2003. doi: 10.1128/JB.185.24.7176-7183.2003.

[23] J. A. Perry, M. B. Jones, S. N. Peterson, D. G. Cvitkovitch, and C. M. Lévesque, “Peptide alarmone signalling triggers an auto-active bacteriocin necessary for genetic competence,” Molecular Microbiology, vol. 72, no. 4, pp. 905–917, 2009, issn: 1365-2958. doi: 10.1111/j.1365-2958.2009.06693.x.

[24] L. Song, W. Wang, G. Conrads, et al.,“Genetic variability of mutans streptococci revealed by wide whole-genome sequencing,” BMC Genomics, vol. 14, no. 1, p. 430, Jun. 2013, issn: 1471-2164. doi: 10.1186/1471-2164-14-430.

[25] V. P. Richards, S. R. Palmer, P. D. Pavinski Bitar, et al., “Phylogenomics and the Dynamic Genome Evolution of the Genus Streptococcus,” Genome Biology and Evolution, vol. 6, no. 4, pp. 741–753, Mar. 2014, issn: 1759-6653. doi: 10.1093/gbe/evu048.

[26] S. Patel and R. S. Gupta, “Robust demarcation of fourteen different species groups within the genus Streptococcus based on genome-based phylogenies and molecular signatures,” Infection, Genetics and Evolution: Journal of Molecular Epidemiology and Evolutionary Genetics in Infectious Diseases, vol. 66, pp. 130–151, Dec. 2018, issn: 1567-7257. doi: 10.1016/j.meegid.2018.09.020.

[27] J. I. Tietz, C. J. Schwalen, P. S. Patel, et al., “A new genome-mining tool redefines the lasso peptide biosynthetic landscape,” Nature chemical biology, vol. 13, no. 5, pp. 470–478, May 2017, issn: 1552-4450. doi: 10.1038/nchembio.2319.

[28] R. W. Mair, D. B. Senadheera, and D. G. Cvitkovitch, “CinA is regulated via ComX to modulate genetic transformation and cell viability in Streptococcus mutans,” Fems Microbiology Letters, vol. 331, no. 1, pp. 44–52, Jun. 2012, issn: 0378-1097. doi: 10.1111/j.1574-6968.2012.02550.x.

[29] S. A. M. Underhill, R. C. Shields, J. R. Kaspar, M. Haider, R. A. Burne, and S. J. Hagen, “Intracellular Signaling by the comRS System in Streptococcus mutans Genetic Competence,” mSphere, vol. 3, no. 5, e00444–18, Oct. 2018, issn: 2379-5042. doi: 10.1128/mSphere.00444-18.

[30] M. Parter, N. Kashtan, and U. Alon, “Facilitated Variation: How Evolution Learns from Past Environments To Generalize to New Environments,” PLOS Computational Biology, vol. 4, no. 11, e1000206, Nov. 2008, issn: 1553-7358. doi: 10.1371/journal.pcbi.1000206.

[31] S. Pollak, S. Omer-Bendori, E. Even-Tov, et al.,“Facultative cheating supports the coexistence of diverse quorum-sensing alleles,” Proceedings of the National Academy of Sciences, vol. 113, no. 8, pp. 2152–2157, Feb. 2016. doi: 10.1073/pnas.1520615113.

[32] N. Aframian and A. Eldar, “A Bacterial Tower of Babel: Quorum-Sensing Signaling Diversity and Its Evolution,” Annual Review of Microbiology, vol. 74, no. 1, pp. 587–606, 2020. doi: 10.1146/annurev-micro-012220-063740.

[33] J. C. Chang, B. LaSarre, J. C. Jimenez, C. Aggarwal, and M. J. Federle, “Two Group A Streptococcal Peptide Pheromones Act through Opposing Rgg Regulators to Control Biofilm Development,” PLoS pathogens, vol. 7, no. 8, M. R. Wessels, Ed., e1002190, Aug. 2011. doi: 10.1371/journal.ppat.1002190.

[34] I. van de Rijn and R. E. Kessler, “Growth Characteristics of Group A Streptococci in a New Chemically Defined Medium.,” Infection and immunity, vol. 27, no. 2, pp. 444–448, Feb. 1980.

[35] C. Engler, R. Kandzia, and S. Marillonnet, “A One Pot, One Step, Precision Cloning Method with High Throughput Capability.,” PloS one, vol. 3, no. 11, H. A. El-Shemy, Ed., e3647, 2008. doi: 10.1371/journal.pone.0003647.

[36] A. B. Shapiro, G. K. Walkup, and T. A. Keating,“Correction for Interference by Test Samples in High-Throughput Assays,” Journal of Biomolecular Screening, vol. 14, no. 8, pp. 1008–1016, Sep. 2009, issn: 1087-0571. doi: 10.1177/1087057109341768.

[37] L. Buitinck, G. Louppe, M. Blondel, et al., “API design for machine learning software: Experiences from the scikit-learn project,” arXiv:1309.0238 [cs], Sep. 2013. arXiv: 1309.0238 [cs].

