## Supplementary Material for "Crosstalk between paralogous ComRS quorum-sensing systems diversifies the competence/bacteriocin networks of oral streptococci"

#### Supplementary information

**Supplementary Table 1. Direct ComRS regulons based on ComR box prediction**

**Supplementary Table 2. List of primers and gBlocks**

**Supplementary Table 3. List of plasmids**

**Supplementary Table 4. List of strains**

**Supplementary Table 5. RODEO analysis of bacteriocin clusters predicted in the ComRS networks**

**Supplementary Text. Reporter construction for ComRS network characterization**

**Supplementary Figure 1. Fluorescent protein reporter selection by P*comS1*$^{SMU}$ response characteristics**

**Supplementary Figure 2. *S. criceti* promoter response to XIP**

**Supplementary Figure 3. Response of (A) P*comX*, (B) P*comS1*, and (C) P*comS2* to XIPs in *S. downei* with either ComRS system deleted.**

**Supplementary Figure 4. Concentration-dependent response of P*comS* and P*comX* to XIP in *S. sobrinus***

**Supplementary Figure 5. Alignment of *comRS* region between *S. downei* and *S. sobrinus*, *S. downei* and *S. criceti* based on encoded protein sequences**

**Supplementary Figure 6. Closeup look at kickstart effects of XIP2 on P*comS1* through ComR2 in *S. mutans*.**

**Supplementary Figure 7. Response of P*comX*, P*comS1*, and P*comS2* to XIP in *S. ratti* and *S. ferus***

**Supplementary Figure 8. Concentration-dependent response of P*comX*, P*comS1*, and P*comS2* to XIP in *S. ratti***

**Supplementary Figure 9. Extended analysis and data on effects of base pair changes in ComR boxes in *S. mutans***

**Supplementary Figure 10. Analysis and summary data on effects of base pair changes in ComR boxes in *S. sobrinus***

**Supplementary Figure 11. Detailed figures for dissected ComRS networks in mutans streptococci, and comparison with *S. sobrinus* ComCDE-BlpCRH network**

### Supplementary Table 1. Direct ComRS regulons based on ComR box prediction

| Species | Locations | ComR box | Notes |
|---|---|---|---|
| ***S. sobrinus*** | ***P_comS*** | aggtgacataaatgtcgttt | *comS* |
| | ***P_comX*** | tagtgacattgatgtcacta | *comX* |
| | ***DLJ51_00230 - 00235* intergenic** | atgcgacattaatgtcaggtgacactaatgtctcta | RiPPs cluster A upstream of *comR*'s |
| | ***DLJ51_00245 - 00250* intergenic** | gggggacattaatgtcagggtgacatcaatgtcacta | |
| | ***P_DLJ51_00255*** | gggtgacattgatgtcgttc | |
| | *P_DLJ51_08380* | aagcgacattcctgtcagacgacatctatgtcgcta | Transmembrane protein, followed by glycosyl transferases, mgtA, small peptides |
| ***S. downei*** | *P_171* | agacgacattgatgtcacctgacatcgatgtcgctt | Thiopeptide RiPPs cluster C |
| | ***P_comS1*** | tagtgacatggatgtcacta | |
| | ***comS1 - NCTC11391_00418* intergenic** | gggtgacatggatgtcaggcgacatccatgtcacta | RiPPs cluster A |
| | ***P_comS2*** | gggtgacatagatgtcaggcgacatccatgtcgcta | |
| | ***P_comX*** | tagtgacatttatgtcacta | |
| | *P_NCTC11391_02073* | aggcgacatctgtgtcaggcgacatgaatgtcgcta | Transmembrane protein, SSO P_DLJ51_08380 homolog, followed by glycosyl transferases,… |
| ***S. criceti*** | ***P_comS*** | tagtgacatgaatgtcactt | |
| | ***P_comX*** | tagtgacatcgatgtcgcta | |
| | *P_STRCR_0572* | gaatgacatcaatgtctttt | With superoxide dismutase domain DUF1033 |
| ***S. mutans*** | ***P_comS1*** | acgggacataaatgtcctgt | |
| | ***P_comS2*** | aggtgacactaatgtcgttg | |
| | ***P_SMU_370*** | tagtgacattaatgtcgtct | RiPPs cluster B |
| | ***P_comX*** | atgggacatttatgtcctgt | |
| | *P_SMU_28* | aggtgacattttgtcacta | SMU_28 gene peptide cleavage/export ABC transporter, partial homolog of cluster A |
| | *P_SMU_440* | attagacatggctgtctatc | 50% ID to 6715 polyketide cyclase,DLJ51_02010 |
| ***S. ratti*** (Notations for assembled genome of ACTC 31377, with same sequence in regions-of-interest to FA1) | ***P_comS1*** | acaggacatagatgtcctgt | |
| | ***P_comS2*** | aggtgacaccaatgtcgttg | |
| | *P_FY406_03900* | acctgacactgatgtcgttt | RiPPs cluster B |
| | ***P_comX*** | acaggacatttatgtcctgt | |
| | *P_FY406_01740* | agccgacattaatgtcgcta | FY406_01740,PRD domain-containing protein, followed by PTS frustose transporters |
| | *P_FY406_00780* (multidrug efflux SMR transporter) | agttgacaagcatgtctgaa | |
| ***S. ferus*** | ***P_comS1*** | acaggacattaatgtcctgt | |
| | ***P_comS2*** | tagtgacattgatgtcatgg | |
| | ***P_comS2*** | ccatgacattgatgtcatgg | *comS2*, RiPPs cluster B |
| | ***P_comX1/2*** | acaggacattaatgtcctgt | |
| | *P_DQL21_RS02855 penA* | ttttgacaaaaatgtcatgg | penA, recR |

| | | | |
|---|---|---|---|
| ***S. macacae*** | *P_comS1* | agaggacatagatgtcctgt | |
| | *P_comS2* | ccatgacataaatgtcatgg | |
| | *P_comS2-B* | aggtgacatttatgtcatgg | *comS2*, RiPPs cluster B |
| | *P_comX* | acaggacatttatgtcctgt | |
| | *P_STRMA_1648* | aggtgacattttgtcacta | post ACP cluster A |

- Bolded genes are the ones for which reporters were constructed. The conserved "gacatnnatgtc" motif in ComR boxes are marked red; disagreeing bases to such are marked green.

### Supplementary Table 2. List of primers and gBlocks

| Function | Primer | Sequence |
|---|---|---|
| mutans ΔcomRS1 | SMU-R1-O-F | tcgcttgagccaagaagaaatcga |
| | SMU-R1-U-F | ctgccttttatgctataatcaagctatgga |
| | O1-SMU-R1-U-R | GAAGGTCTCTCTATACCTTTTCCTATAATCTCTGTCTAAACTTTTACT |
| deleting *comS* together | O2-SMU-R1S-D-F | GAAGGTCTCTAATCagacagcccttatgtcagatgatgattt |
| | SMU-R1S-D-R | tggtgatgatgctatcaacgcga |
| | SMU-R1-O-R | acctctacaagtaaaggaattacagcgga |
| R1 cloning | O1-SMU-R1-F | GAAGGTCTCTATAGtcgtttgctgcaagactacgct |
| | O4-SMU-R1-R | GAAGGTCTCTAGTCttatgtcccgttctgagaatctttttgcca |
| mutans ΔcomRS2 | SMU-R2-O-F | tgataccgctgagtattatggacatgg |
| | SMU-R2-U-F | tctgccaaaacaattgcacagact |
| | O1-SMU-R2-U-R | GAAGGTCTCTCTATgatggctcagttcctctcaatatgct |
| | O2-SMU-R2S-D-F | GAAGGTCTCTAATCgccgtgcgctaataatgaacatcta |
| | SMU-R2-D-R | agagtcggctaatattgttctggattact |
| | SMU-R2-O-R | taacggcattacgtcttttggct |
| | TS-O1-SMU-R2-F | GAAGGTCTCTATAGacccactacagatattatgaactgcacca |
| | TS-O4-SMU-R2-R | GAAGGTCTCTAGTCtgctagtctatcccatcatttgccatct |
| downei ΔcomRS1 | dow-R1-O-F | actataccatcaaacaggtccgca |
| | TS-dow-R1-U-F | acgtcccttactatatgctttgtgagt |
| | TS-O1-dow-R1-U-R | GAAGGTCTCTCTATgaggttctccttcatagattattgagccct |
| | O2-dow-R1S-D-F | GAAGGTCTCTAATCtaagaagaggcctaagtaaggataactcaaaga |
| | dow-R1-D-R | tcctttcgctatattaatttatcaataaagtaggca |
| | dow-R1-O-R | actggttaagtcagatttcataccatcca |
| | TS-O1-dow-R1-F | GAAGGTCTCTATAGagatataagtctgttattctggcgacaca |
| | TS-O4-dow-R1-R | GAAGGTCTCTAGTCctatgccaggttatccctatccttttct |
| downei ΔcomRS2 | dow-R2-O-F | tcaaaatcaagggtcagaagaaaagcct |
| | dow-R2-U-F | acctccaatactcaacgagcagacat |
| | O1-dow-R2-U-R | GAAGGTCTCTCTATtacaagtacccccctgagtttaaatctaacttaatt |
| | TS-O2-dow-R2S-D-F | GAAGGTCTCTAATCacagtattgtttcggtagtatagataaatagaaagga |
| | dow-R2-D-R | agacctcaagctattcacagaactattaga |
| | dow-R2-O-R | agtgctatgctacgaaaagtagcaaaa |
| | O1-dow-R2-F | GAAGGTCTCTATAGtcctcaaaaggggtgagtcttttgt |
| | O4-dow-R2-R | GAAGGTCTCTAGTCagcaaaggaatctattcaatcccatcaatt |
| criceti ΔcomRS | cri-R-O-F | tcttaacgggatagctgatgttgttgt |
| | TS-cri-R-U-F | tccgctatcaaaaatggcggga |
| | O1-cri-R-U-R | GAAGGTCTCTCTATaaaggccctccttctagcttttattacc |
| | O2-cri-RS-D-F | GAAGGTCTCTAATCtaggtgaacgcattagcaaaattaaagaga |
| | TS-cri-R-D-R | tcctaactcgtatttctatctgcttagcca |
| | cri-R-O-R | acctgaactaaacgcaactttttcaaact |
| | O1-cri-R-F | GAAGGTCTCTATAGtcaattttttgggttgattctactcccct |
| | O4-cri-R-R | GAAGGTCTCTAGTCtcactagactagatcatcctcagcttttt |
| ferus ΔcomRS1 | fer-R1-O-F | agactttctcgatggtttggccaa |
| | fer-R1-U-F | ttggagacctatcaggatcagcca |
| | O1-cri-R1-U-R | GAAGGTCTCTCTATtccattttcccaatcttgttattactaatagtttact |
| | TS-fer-RS1-D-F | GAAGGTCTCTAATCtaagctacaagaattcggagaacactaattga |
| | fer-R1-D-R | tccaatccttctcgatcgacatcca |
| | fer-R1-O-R | acgctcctctgcaatattaaccga |
| | O1-fer-R1-F | GAAGGTCTCTATAGtggggttaatatgacttgcgtcaact |

| | | |
|---|---|---|
| | O4-fer-R1-R | GAAGGTCTCTAGTCtcctgttattttttaaatcttgtgcccact |
| ferus ΔcomRS2 | fer-R2-O-F | tgttatgcttttcgtatcattgagcgt |
| | fer-R2-U-F | tatcgcccacttaacaagcaagga |
| | O1-fer-R2-U-R | GAAGGTCTCTCTATaaaatccccacctttatactaccattctaacata |
| | O2-fer-R2S-D-F | GAAGGTCTCTAATCttctaattagtctgttgcttgacggattaat |
| | fer-R2-D-R | aaaatggcaggcaaattgctagga |
| | fer-R2-O-R | actaaggttgatgcaagatattcatatcct |
| | O1-fer-R2-F | GAAGGTCTCTATAGagccctaagcgatgtaagctgt |
| | O4-fer-R2-R | GAAGGTCTCTAGTCttaaatatgatcagcttttctttcgtctctga |
| ratti ΔcomRS1 | rat-R1-O-F | tgggttcactttgggctgacct |
| | rat-R1-U-F | accatcatgatgggacgtaccca |
| | O1-rat-R1-U-R | GAAGGTCTCTCTATtctctgatatgtcagctgatttttattcatttga |
| | O2-rat-RS1-D-F | GAAGGTCTCTAATCatctagtttgtgaacactaaaacttccaga |
| | rat-R1-D-R | tctccccagcaccaatcataatgt |
| | rat-R1-O-R | agtattccatgtgccctgtaatacca |
| | O1-rat-R1-F | GAAGGTCTCTATAGtccggtttattacactaaacgcgtagat |
| | O4-rat-R1-R | GAAGGTCTCTAGTCaggacatctatgtcctgttttggtagt |
| ratti ΔcomRS2 | rat-R2-O-F | agagcaagggcgaaaggtctgt |
| | rat-R2-U-F | agcggcaataatttgccaacaagga |
| | O1-rat-R2-U-R | GAAGGTCTCTCTATctcccctagtctttttgtatgttcattatacc |
| | O2-rat-R2S-D-F | GAAGGTCTCTAATCatggcaatatactaataattaatattgattaggaacaact |
| | rat-R2-D-R | tgctagtattaaccctactactatcaattttaggaaga |
| | rat-R2-O-R | aggtgtggctattctcacctaaagga |
| | TS-O1-rat-R2-F | GAAGGTCTCTATAGagcaccttctccattaggcgcta |
| | TS-O4-rat-R2-R | GAAGGTCTCTAGTCagtgtctcctgctagtctatcccat |
| criceti P*comX* | cri-P*comX*-F | TACggtctcgATAGacggataaatagatataaggtcgctgtggga |
| | cri-P*comX*-R | GCTggtctccataaaatctccttcaatttcttttgcttaattataataaga |
| | criTTAT-mNGFP-F | TACggtctcgttatATGGTGTCGAAAGGAGAGGAGGATAAT |
| replacement for SMU P*comX* | pRW17-TCTA-R2 | GCTGGTCTCCTAGAtgtagtccctcactgattaagcattggta |
| | pRW17-seq-F | acgttaagggattttggtcatgagatt |
| ***comX* KO** | 67*comX*-o-F | cataacgttaagttagtaagagccctgaga |
| 6715 | 67*comX*-U-F | tactaatcgctcgaggacttgacca |
| | O1-67*comX*-U-R | GAAGGTCTCTCTATctaaattctcctttaaattttttcttaaattataataggactctg |
| | O2-67*comX*-D-F | GAAGGTCTCTAATCcaactaagttcacttcctacaagccg |
| | 67*comX*-D-R | aaccatacgagcgtcacgagtaa |
| | 67*comX*-o-R | acagttggcctcaaaggtaaaacgt |
| | SMU*comX*-o-F | cataaataatgaagcatctttacctaggtgct |
| SMU | SMU*comX*-U-F | tcaattgtattgagttgaatcgggtagca |
| | O1-SMU*comX*-U-R | GAAGGTCTCTCTATtcaaaatcttcttccatctattacgatgacct |
| | O2-SMU*comX*-D-F | GAAGGTCTCTAATCaagtgattttaaggaaaaatagttaaaaaggggaaaga |
| | SMU*comX*-D-R | aaaaccgcaaatcatgacgttcatt |
| | SMU*comX*-o-R | tgccaaaattaaccacattctaagtaacttt |
| | DOW*comX*-o-F | tcaggagaatcgagttggttctct |
| downei | DOW*comX*-U-F | tcaatcttggcttgaatcagtcctagt |
| | O1-DOW*comX*-U-R | GAAGGTCTCTCTATttcttcgtccattcaatgtctcctttt |
| | O2-DOW*comX*-D-F | GAAGGTCTCTAATCcgatgaccactgaagctaacctttt |
| | DOW*comX*-D-R | tgctcaccaaggctgtcaaaag |
| | DOW*comX*-o-R | tcgctcgacctgttcttcact |
| criceti | CRIU*comX*-check-R | tcccacagcgaccttatatctatttatccgt |
| | CRI*comX*-U-F | tgttagaatctcaattctcaattgaggttct |

| | | |
|---|---|---|
| | O1-CRI*comX*-U-R | GAAGGTCTCTCTATatctttgagtttttctttatccatataaaatctccttca |
| | O2-CRI*comX*-D-F | GAAGGTCTCTAATCttttgacaacgatgattagaagagatatccat |
| | CRI*comX*-D-R | aagataagctctcaaataggggaggtcta |
| | CRI*comX*-o-R | tgaaagtttgaggccgtagcgt |
| ratti | RAT*comX*-U-F | agatgaatattctaaaacacctcattgaggt |
| | O1-RAT*comX*-U-R | GAAGGTCTCTCTATatagggcattttgttaaaacgacgttt |
| | O2-RAT*comX*-D-F | GAAGGTCTCTAATCttttgatgacaccagtttaagggaaca |
| | RAT*comX*-D-R | tctcaccttgtaaacttccctgca |
| | RAT*comX*-o-R | agctcgtgctgattttcagcct |
| ferus *comX*1 | FER*comX*-U-F | ttgagacagcacctgagggt |
| | O1-FER*comX*-U-R | GAAGGTCTCTCTATaatctcttcataagccatctgatgaaact |
| | O2-FER*comX*1-D-F | GAAGGTCTCTAATCacttcagggaatgaaaggatttagctgt |
| | FER*comX*1-D-R | aatagagccccatttcatatctgcct |
| | FER*comX*1-o-R | tccatatcaaaaataactgcctgtcgt |
| ferus *comX*2 | O2-FER*comX*2-D-F | GAAGGTCTCTAATCaagacttcagggaataaaaaggatttagct |
| | FER*comX*2-D-R | tcagattttcatttgccaagagcatca |
| | FER*comX*2-o-R | tttcaagcatcccctgctgaact |
| **P*comS* fusions** | | |
| 6715 | 67P*comS*-F | GAACggtctctATAGtaaagtttctacggagcttcattgtatagaga |
| | 67P*comS*-R | GAAGGTCTCTtattaaactccctttcattttataaacctattctaca |
| | 67TATT-mNG-F | GAAggtctctAATAATGGTGTCGAAAGGAGAGGAGGATAAT |
| | 67TATT-mSC-F | GAAggtctctAATAATGGTAAGTAAAGGCGAAGCAGTCA |
| | mSC-O4-R | GAAggtctcTagtcTTACTTATATAACTCATCCATTCCACCAGTAGAGT |
| downei-S1 | dowP*comS1*-F | GAAggtctctATAGaattttgcaaccttgctaggggatca |
| | dowP*comS1*-R | GAAGGTCTCTtttctccagcctttcttttgatggtta |
| | dows1-mNG-F | GAAggtctctgaaaATGGTGTCGAAAGGAGAGGAGGATAAT |
| | dows1-mSC-F | GAAggtctctgaaaATGGTAAGTAAAGGCGAAGCAGTCA |
| downei-S2 | dowP*comS2*-F | GAAggtctctATAGggtaaggcgggatcctagtagct |
| | dowP*comS2*-R | GAAGGTCTCTttggattaaaaacttcctttcatttgataagtctattct |
| | dows2-mNG-F | GAAggtctctccaaATGGTGTCGAAAGGAGAGGAGGATAAT |
| | dows2-mSC-F | GAAggtctctccaaATGGTAAGTAAAGGCGAAGCAGTCA |
| criceti-S | criP*comS*-F | GAAggtctctATAGaacttcgcaaccttgttgggaga |
| | criP*comS*-R | GAAGGTCTCTtttctctatcctttcttttcgatattttcattatatataaaatcttca |
| mutans-S1 | smuP*comS1*-F | GAAggtctctATAGtgcccagttaacagaaaacattaatttaagaga |
| | smuP*comS1*-R | GAAGGTCTCTcctgttattctcctttctttttgatatcattcat |
| | smus1-mNG-F | GAAggtctctCAGGATGGTGTCGAAAGGAGAGGAGGATAAT |
| | smus1-mSC-F | GAAggtctctCAGGATGGTAAGTAAAGGCGAAGCAGTCA |
| mutans-S2 | smuP*comS2*-F | GAAggtctctATAGtttcaaagtaaaaggggaagtcactagtaca |
| | smuP*comS2*-R | GAAGGTCTCTttgataaagacttcctttcatttgatgagtctatt |
| | smus2-mNG-F | GAAggtctctTCAAATGGTGTCGAAAGGAGAGGAGGATAAT |
| | smus2-mSC-F | GAAggtctctTCAAATGGTAAGTAAAGGCGAAGCAGTCA |
| ratti-S1 | ratP*comS1*-F | GAAggtctctATAGgtttgcgaaattaactgaaaataccacattaaga |
| | ratP*comS1*-R | GAAGGTCTCTatccttcttctcctttctttttgatgttattc |
| | rats1-mNG-F | GAAggtctctGGATATGGTGTCGAAAGGAGAGGAGGATAAT |
| | rats1-mSC-F | GAAggtctctGGATATGGTAAGTAAAGGCGAAGCAGTCA |

| | | |
|---|---|---|
| ratti-S2 | ratP*comS2*-F | GAAggtctctATAGagtggggtaaaattactagtctaaaaagatgatt |
| | ratP*comS2*-R | GAAGGTCTCTtgataaagacttcctttcatttgataagtctattcta |
| ferus-S1 | ferP*comS1*-F | GAAggtctctATAGgctcaattaattggaaatgcttatttggca |
| | ferP*comS1*-R | GAAGGTCTCTtgctttctcctttctcttattttatggctttatt |
| | fers1-mNG-F | GAAggtctctAGCAATGGTGTCGAAAGGAGAGGAGGATAAT |
| | fers1-mSC-F | GAAggtctctAGCAATGGTAAGTAAAGGCGAAGCAGTCA |
| ferus-S2 | ferP*comS2*-F | GAAggtctctATAGtggagataatattttagaagctaatctcagagacga |
| | ferP*comS2*-R | GAAGGTCTCTtttctatcatcctttcacatttacgattttattataagt |
| **Cloning all FP to UP*comS*** | mSC-check-R | TGCGGACTAAGGATATCCCAGCT |
| mTB2 | SMUS1-mTB2-F | GAAggtctctCAGGATGGTCAGCAAGGGAGAAGAATTGATCA |
| | mTB2-O4-R | GAAggtctcTagtcTTAGTTCAACTTGTGACCCAATTTAGATGGT |
| | mTB2-check-R | ACCTTCATAGGGTTTACCCTCACCT |
| sfGFP | SMUS1-sfGFP-F | GAAggtctctCAGGATGAGCAAAGGAGAAGAACTTTTCACTGGA |
| | sfGFP-O4-R | GAAggtctcTagtcTTAGTGGTGGTGGTGGTGGT |
| | sfGFP-check-R | TGTAGCATCACCTTCACCCTCTCCA |
| mTFP1 | SMUS1-mTFP1-F | GAAggtctctCAGGATGGTATCTAAAGGCGAAGAAACGACTAT |
| | mTFP1-O4-R | GAAggtctcTagtcTTACTTGTACAACTCGTCCATTCCGT |
| | mTFP1-check-R | TCGTATGGCTTTCCCTCACCTTCT |
| mTq2 | SMUS1-mTq2-F | GAAggtctctCAGGATGGTTTCTAAAGGTGAGGAATTATTTACTGGT |
| | mTq2-O4-R | GAAggtctcTagtcTTATTTATACAACTCGTCCATTCCAAGTGT |
| | mTq2-check-R | AGCTTTCCATACGTAGCATCTCCT |
| ***comS* KO** | | |
| suitable for reporter int | O1-p17-int-F | GAAGGTCTCTATAGagatcctttgatcttttctacggggtct |
| 6715 *comS* | 67*comS*-O-F | tactttttcagaactaaacgaggaagactt |
| | 67*comS*-U-F | ggtggtatccacgtattggaattcca |
| | O1-67*comS*-U-R | GAAGGTCTCTCTATaggtgatctcttaaaagaaatcactcttctct |
| dowS1 | dowS1 -O-F | tcgctatggagattgccgttcga |
| | dowS1 -U-F | tcattgaagcagccaacgggt |
| | O1-dowS1 -U-R | GAAGGTCTCTCTATagaaaaggatagggataacctggcatag |
| dowS2 | dowS2-O-F | agacctatctaaagaacaggtttggagt |
| | dowS2-U-F | tcctcaaaaggggtgagtcttttgt |
| | O1-dowS2-U-R | GAAGGTCTCTCTATagcaaaggaatctattcaatcccatcaatt |
| criS | criS-O-F | atcaattgtgcgaggatgggagt |
| | criS-U-F | tcaatttttgggttgattctactcccct |
| | O1-criS-U-R | GAAGGTCTCTCTATtcactagactagatcatcctcagcttttt |
| US1 | US1-O-F | tcgcttttaataagctagcgcttattca |
| | US1-U-F | tcgtttgctgcaagactacgct |
| | O1-US1-U-R | GAAGGTCTCTCTATttatgtcccgttctgagaatctttttgcca |
| US2 | US2-O-F | aatcaggaaaacagcaatctgatcct |
| | US2-U-F | acccactacagatattatgaactgcacca |
| | O1-US2-U-R | GAAGGTCTCTCTATtgctagtctatcccatcatttgccatct |
| fS1 | fS1-O-F | tccttggagagtttttctggctct |
| | fS1-U-F | tggggttaatatgacttgcgtcaact |
| | O1-fS1-U-R | GAAGGTCTCTCTATtcctgttattttttaaatcttgtgcccact |

| | | |
|---|---|---|
| fS2 | fS2-O-F | aaaagggcttcagttggcagtt |
| | fS2-U-F | agccctaagcgatgtaagctgt |
| | O1-fS2-U-R | GAAGGTCTCTCTATttaaatatgatcagcttttctttcgtctctga |
| rS1 | rS1-O-F | tcaaaccgctgctggaagca |
| | rS1-U-F | tccggtttattacactaaacgcgtagat |
| | O1-rS1-U-R | GAAGGTCTCTCTATtctatgtcctgttttggtagtccttct |
| rS2 | rS2-O-F | agctgacacaaagcttgttgtcat |
| | rS2-U-F | agcaccttctccattaggcgcta |
| | O1-rS2-U-R | GAAGGTCTCTCTATagtgtctcctgctagtctatcccat |
| 6715 PcomR1 reporter | 67PR1-F | GAAggtctctATAGGCAGCTTGATCTGAGAAAATCAGGCT |
| | 67PR1-R | GAAGGTCTCTACTGGGCCCTCCAATTTAGTCCT |
| | 67PR1-mNG-F | GAAggtctctCAGTATGGTGTCGAAAGGAGAGGAGGATAAT |
| 6715 PcomR2 reporter | 67PR2-F | GAAGGTCTCTATAGtccttatagataagacaagcactctcaacaac |
| | 67PR2-R | GAAGGTCTCTaatcgtcctccaattcttggttcatt |
| | 67PR2-mNG-F | GAAggtctctgattATGGTGTCGAAAGGAGAGGAGGATAAT |
| 6715 04555 reporter | 67P4555-F | GAAggtctctATAGtcattgtaggctatctgcttggacttt |
| | 67P4555-R | GAAGGTCTCTagataactctcctttttcgtcaaaaaccct |
| | 67P4555-mNG-F | GAAggtctctatctATGGTGTCGAAAGGAGAGGAGGATAAT |
| 6715 04575 reporter | 67P4575-F | GAAggtctctATAGtctagttagtggcaacgccccgtt |
| | 67P4575-R | GAAGGTCTCTaatacctccattcctgttaagagaagttaaaaact |
| | 67P4575-mNG-F | GAAggtctcttattATGGTGTCGAAAGGAGAGGAGGATAAT |
| 6715 00231 reporter | 67P231-F | GAAggtctctATAGagttaagctcttaacccgagtagggt |
| | 67P231-R | GAAGGTCTCTgcttatcccctttcagttataatttaatttcgaaattagt |
| | 67P231-mNG-F | GAAggtctctaagcATGGTGTCGAAAGGAGAGGAGGATAAT |
| new primers for DowS1-U | dowS1-U-R2 | GAAGGTCTCTCTATcatgtcactatgccaggttatccct |
| **SMU bcin1 reporter** | muP370-F | GAAggtctctATAGactttaatttagctccatttttacagcacct |
| | muP370-R | GAAggtctctacgttttcctttcttaacggtatagcat |
| | muP370-mNG-F | GAAggtctctacgtATGGTGTCGAAAGGAGAGGAGGATAAT |
| | muP370-int-check-F | actgggttctatctttgttagtaatgtga |
| | muP370-int-check-R | gcatcaagtgcaacagccttgt |
| **gBlocks for R'-P*comS* reporters** | | |
| | 6715 R-P*comS* | ACTGGTCTCTaTagtaagtaaaGCAGCTTGATCTGAGAAAATCAGGCTttttttttgctttttgaaagcgaccaagtgcaaattcattgcttgatatgtcattaaactgttattttttgatattgcacagatatagaatagaaaggtataattaacagAGGACTAAATTGGAGGGCCCAGTatgacaataaaagcagatttgggacgacggattaaggccgaaagaaaccggaaaaaaatatcgcagagttcattttgcgggctgggagatattctagatgatcccaccattattgctgatgttaagatggaaatgaaaaatgatgggatagatgaataaagtttctacggagcttcattgtatagagaagagtgatttcttttaagagatcacctttttagtttttggaaaaggtgacataaatgtcgttttgttttgagctgaatgggtgtagaataggtttataaaatgaaagggagtttaataAGAGACCTTC |
| | downei R1-P*comS1* | ACTGGTCTCTaTagagatataagtctgttattctggcgacacaaagttttgctagaatgggggtgagagatgaagggatgaaagtagtcattgaagcagccaacgggttcttgaaaagtaagttttctggagttaatagctttgaaagtcttacgttttctgacttgttaagggctcaataatctatgaaggagaacctcatgcctttcaaatcaaaaatcggaccccagattcgtcaaaa |

| | | |
|---|---|---|
| | | gcgggaggccaaacagctctctcgggaaaagctctgccatgatggtagtcgcttaacgcagctttatgatcaagccattaattttgcaaccttgctaggggatcaggtcttggctgatggcctacgtgaagaaaaggatagggataacctggcatagtgacatggatgtcactatctttttggtggaagtaatatataatgtaaccatcaaaaagaaaggctggagaaaAGAGACCTTC |
| | downei R2-P*comS2* | ACTGGTCTCTaTagggtgagtcttttgtgggatgttcagcttatatcagttattcctatctagtgatgacattgctgtcacaagattgacattcttttgatattgtacagagatatcagaaaagggtataattaagttagatttaaactcaggggggtacttgtagtggctatgaaggaagaattgggacaactcatcaagaaaaaaagggaagcgaagaagctttcgcagaagaacttatgtggcatgttgggagagatactgggcgatcaagtaataattaatgatatcaaaaaagaaatgaaaattgatgggattgaatagattcctttgctagtagtgggtaaggcgggatcctagtagctttgtatttagggctactatcatttagaaaagggtgacatagatgtcaggcgacatccatgtcgctatttttttaggagagaaataggtagaatagacttatcaaatgaaaggaagtttttaatccaaGAGACCTTC |
| | criceti R-P*comS* | ACTGGTCTCTaTagttgattctactccccttttttgtttgctgggaaaaaggaaattttaatctttgtttaagttttagaatcttaactttttctgtaaataaagatacaagattgttatttaaacgatactaatatttgataaaatagagatgagagatgaagggtaataaaagctagaaggagggcctttatgcctttcaaatcaacaattggtcccagaattcgtcagactagagaaagccgaaatctctctcgagactcagtctgtcgcgatggtagtcatctgactcaactttatgatcaagccattaacttcgcaaccttgttgggagatgaggttttagcagatggtctgcgtgatgaaaaagctgaggatgatctagtctagtgacatgaatgtcactttctttttggtgaagattttatataatgaaaatatcgaaaagaaaggatagagaaaAGAGACCTTC |
| | SMU R1-P*comS1* | ACTGGTCTCTaTaggattaatcgtttgctgcaagactacgcttatgtcttatgcttcctttttgaaactcgctgaatgtgttcccaaaagatgtcagaaaaatatatagaagtctagtaaatttactatattatagagttatgataaaatagtaaaagtttagacagagattataggaaaaggttttatgttaaaagattttgggaagaaaattaagagcctgaggttagaaaaggggctgaccaaagaagctgtctgccttgatgaagcccagttaacagaaaacattaatttaagagaaaatttggagaaagaatggcaaaaagattctcagaacgggacataaatgtcctgttctttttttgaaggatcatttataatgaatgatatcaaaaagaaaggagaataacaggAGAGACCTTC |
| | SMU R2-P*comS2* | ACTGGTCTCTaTagacccactacagatattatgaactgcaccatagctgataactaaacttaaagacagacatagcaacaagtggaaaatctaaaccgaaacatttcagaatccagaataacagaaaaccattcttgatactgtacagacataggagataatgatataatcagcatattgagaggaactgagccatcatgacaataaaagaagatttaggacggcggattaaagcagaaaggaatcgcaagcaactcacacaaagccttttgtgcggagatatgggcgaagttctaggtgatcctgttataattgcagatgttaagatggagatggcaaatgatgggatagactagcaagagaggttattaagtttagcataacaaagaactttagcgtctatactttcaaagtaaaaggggaagtcactagtacaaaaaggatgattctctttagaaaaatattctttttccaagaggtgacactaatgtcgttgttgctttgaggaatctggggtagaatagactcatcaaatgaaaggaagtctttatcaaAGAGACCTTC |
| | ferus R1-P*comS1* | ACTGGTCTCTaTagtaggccttatttattttggggttaatatgacttgcgtcaactttaacggtatatcaaaaggtttttgtaaagtaaagttacttgatagatttttttgtttcaactgttcagatcttgtagcttaacaagaaagcgtcataaaaaaacttgttaattttaatataattaagtaaactattagtaataacaagattgggaaaatggatgttaaagaattttgggattaaagtcagagagctgcgagcggaaaaagggctaaccaaagaaatcttctgccaagacgaaacggaactgtctatccgggaaagctatcattctgctgtgatgcttgctcaattaattggaaatgcttatttggcagaaaaattagaagaagagtgggcacaagatttaaaaaaataGcaggacattaatgtcctgttattttttttatgctcttttctataataaagccataaaataagagaaaggagaaagcaAGAGACCTTC |
| | ferus R2-P*comS2* | ACTGGTCTCTaTaggaagcgggtcaggggaggaatccagcagccctaagcgatgtaagctgtgtgctctttttttatggctgtttaaagcaccatatatgaggaaacgtgcaatctgtaactattaaaacgttttcaattcttttatgtttcccagcctgttttatgttagaatggtagtataaaggtggggattttatgagtatcaaggaaacaattgggaaaaaaatcagggaagaaagagagaagaagggattctctcgagagcagttgtgtgggggcgaggaagaactgactaaagaatgttatgatttagcgaccttattagcacaaaattttggagataatattttagaagctaatctc |

| | | |
|---|---|---|
| | | agagacgaaagaaaagctgatcatatttaaataacgtagaaagaagaccatgacattgat gtcatggtcttctttttactttttctataatgaagccataaagtttgaaagtgattcatattgat gcaagactttcaactttgggcagtcagtaagtcctcagtagtgacattgatgtcatggtcaa atatggctgccagacttataataaaatcgtaaaatgtgaaaggatgatagaaaAGAG ACCTTC |
| | ratti R1-P*comS1* | ACTGGTCTCTaTagtccggtttattacactaaacgcgtagatgagattttcaaac gtatcggtttggaagattaacagactgcagaaagctgtgacaacagtcgcagtttttttgc gtttacggattcaaatgaataaaaatcagctgacatatcagagaaatttactatattatagac ttatgatagaataataaaaaatgagaaagagattttatgttagaagattttggggagagagt aaaacgtctgcggttagaaaagggactgaccaaagaagttttctgtcaagatgagtctga gctgtctatgcggtatgactcggctattttgtttgcgaaattaactgaaaataccacattaag agaaaagttagaaagagagtggcagaaggactaccaaaacaggacatagatgtcctgtt cttttcttctccgatattttataatgaataacatcaaaaaagaaaggagaagaaggatAG AGACCTTC |
| | ratti R2-P*comS2* | ACTGGTCTCTaTagagcaccttctccattaggcgctagtaactgtatttcagaag ctgtactagggtggggcaaagtgaaggcaagaacctatagttttgatttttgaagagtaga atcgtcatcaaattgtccctttcataatattgcacaatctaggggagaacggtataatgaac atacaaaaagactaggggagacTatcatgacaataaaagaagatttgggacagcggat taaagacaaaagaaatcaacagcaactgacacaaagccttttgtgcggagacgaaaca aagctgaccgaactctaccagcaagccttggtgatgggagaaatcttaggcgactctatt atcattgcagatgtcaagatggagatggcaaatgatgggatagactagcaggagacact attaagcttttagttacaacaaatatctttagcttctgtattctcaaagcagtggggtaaaatta ctagtctaaaaagatgattcatttttagtagtttaggtatcatcactttttttacaagaggtgaca ccaatgtcgttgctgctttgaggaatatggagtagaatagacttatcaaatgaaaggaagt ctttatcaaAGAGACCTTC |
| **Primers for gBlocks** | | |
| | 67PS gB-F | ACTGGTCTCTaTagtaagtaaaGCAGCT |
| | 67PS gB-R | GAAGGTCTCTtattaaactccctttcattttataaac |
| | dowPS1 gB-F | ACTGGTCTCTaTagagatataagtctgttattct |
| | dowPS1 gB-R | GAAGGTCTCTtttctccagccttt |
| | dowPS2 gB-F | ACTGGTCTCTaTagggtgagtcttttg |
| | dowPS2 gB-R | GAAGGTCTCttggattaaaaacttcctttc |
| | criPS gB-F | ACTGGTCTCTaTagttgattctactccc |
| | criPS gB-R | GAAGGTCTCTtttctctatcctttcttttcg |
| | SMU PS1 gB-F | ACTGGTCTCTaTaggattaatcgtttgct |
| | SMU PS1 gB-R | GAAGGTCTCTcctgttattctcctttct |
| | SMUPS2 gB-F | ACTGGTCTCTaTagacccactacagata |
| | SMUPS2 gB-R | GAAGGTCTCTtgataaagacttcctttca |
| | ferPS1 gB-F | ACTGGTCTCTaTagtaggccttatttattttg |
| | ferPS1 gB-R | GAAGGTCTCTtgctttctcctttctc |
| | ferPS2 gB-F | ACTGGTCTCTaTaggaagcgggt |
| | ferPS2 gB-R | GAAGGTCTCTtttctatcatcctttcacatt |
| | ratPS1 gB-F | ACTGGTCTCTaTagtccggtttattaca |
| | ratPS1 gB-R | GAAGGTCTCTatccttcttctcctttct |
| | ratPS2 gB-F | ACTGGTCTCTaTagagcaccttctc |
| | ratPS2 gB-R | GAAGGTCTCTtgataaagacttcctttca |
| **All reporters with mNG** | | |
| 6715 | 67P*cinA*-F | GAAggtctcTATAGctgcatagggtcaaaatacgctttct |
| | 67P*cinA*-R | GAAggtctcTCCATtaggtcacccacttatctattcgtaaagtca |
| | P4575-F2 | GAAggtctcTATAGagacaggaaaatttcatccatgtcgt |
| | P4575-R | GAAggtctcTCCATaatacctccattcctgttaagagaagttaaaaact |
| Downei | dowPR1-F | GAAggtctcTATAGtcgctatggagattgccgttcga |

| | | |
|---|---|---|
| | dowPR1-R | GAAggtctcTCCATgaggttctccttcatagattattgagccct |
| Bcin1 | P417-F | GAAggtctcTATAGgcgattttggtcccgctctct |
| | P417-R | GAAggtctcTCCATcagaatctcctttctatcaaatttaattctagaattact |
| | dowPR2-F | GAAggtctcTATAGtcctcaaaaggggtgagtcttttgt |
| | dowPR2-R | GAAggtctcTCCATtacaagtaccccctgagtttaaatctaacttaatt |
| Bcin3 | P171-F | GAAggtctcTATAGacgtggcatcatcgagttagacga |
| | P171-R | GAAggtctcTCCATctttcattttatttctcctttctttcaaatttcattct |
| | PfasA2-F | GAAggtctcTATAGagattgacatgggattggccct |
| | PfasA2-R | GAAggtctcTCCATagataactctcctttttcgtcaaaaaccct |
| Bcin2 | PubaA-F | GAAggtctcTATAGaaaaactcctttaaagacttaatcagagcca |
| | PubaA-R | GAAggtctcTCCATaataatataattcctccgtttcttgtaaaaagaagtt |
| ==*cinA* | PyfaY-F | GAAggtctcTATAGcacggacaaagcaagtctattattggta |
| | PyfaY-R | GAAggtctcTCCATtaggtcacctctacttatctattcgtaaagtca |
| criceti | PsakP2-F | GAAggtctcTATAGagtcatctggaattgaacagccaga |
| | PsakP2-R | GAAggtctcTCCATtttgctaccttcttaaacattgttattaaactagct |
| | criP*cinA*-F | GAAggtctcTATAGtgctctaacaatacagattttcctccct |
| | criP*cinA*-R | GAAggtctcTCCATtatatatggtcacctacttatctattcgtaaagtca |
| SMU | | |
| | P*nlmA*-F | GAAggtctcTATAGaccaccagttcacactggtggt |
| | P*nlmA*-R | GAAggtctcTCCATatgataaacacccctttttcattttaaatattgtct |
| | uP*cinA*-F | GAAggtctcTATAGacaggcagcaggtttaattaatgatca |
| | uP*cinA*-R | GAAggtctcTCCATtctaacctcctcaatattatattcgtaaaagattga |
| | muP370-F | GAAggtctcTATAGactttaatttagctccatttttacagcacct |
| | muP370-R | GAAggtctcTCCATacgttttcctttcttaacggtatagcat |
| | uPcomE-F | GAAggtctcTATAGactaccaagtgctattactgtcgtca |
| | uPcomE-R | GAAggtctcTCCATtatttctcctttaatcttctatttaggttagctga |
| **SMU bcin deletions** | | |
| 370 cluster | mu370-O-F | tctggaattgagtgagttgttcgga |
| | mu370-U-F | tgcataaatagggatactggcttgct |
| | O1-mu370-U-R | GAA**GGTCTC**TCTATacgttttcctttcttaacggtatagcat |
| | O2-379-D-F | GAA**GGTCTC**TAATCtgctgctgttctgacatagacagat |
| | 379-D-R | acagcggactatgatcagacaact |
| | 379-O-R | tgtcagagaatatcctagacagttttcatga |
| | O1-370-F | GAAGGTCTCTATAGactttaatttagctccatttttacagcacct |
| | O4-379-R | GAAGGTCTCTAGTCacctgctaagggtattacagggct |
| comDE | comE-O-F | aggcgactctagctaaacagcat |
| | comE-U-F | acggcaaaaagttgctgacggt |
| | O1-comE-U-R | GAA**GGTCTC**TCTATactgttaaccactttgactcacctatgct |
| | O2-comD-D-F | GAA**GGTCTC**TAATCtctcatccacgacagcacactt |
| | comD-D-R | tgtagcagccgaactttcttgtgaa |
| | comD-O-R | tgattctgctggcaaattcgctt |
| | O1-comE-F | GAAGGTCTCTATAGagcataggtgagtcaaagtggttaacagt |
| | O4-comD-R | GAAGGTCTCTAGTCaactatctcttagagaataggcctcctct |
| kanR check primers | aph3-i-F | TATCGGGGAAGAACAGTATGTCGA |
| | aph3-i-R | CTCCCACCAGCTTATATACCTTAGCA |
| **Promoter variant gBlocks** | muP*comX*-V1 | GAACggtctctATAGaactgtttaatctgttagtcagaggttgtaaataaaacagcc agttaagatgTgacatttatgtcctgttcttaaagtctttttcgttttataataattttattataaa aggaggtcatcgtaatagATGGAgagaccTTC |
| | muP*comX*-V2 | GAACggtctctATAGaactgtttaatctgttagtcagaggttgtaaataaaacagcc agttaagatgggacatttatgtcGtgttcttaaagtctttttcgttttataataattttattataaa aggaggtcatcgtaatagATGGAgagaccTTC |

| | | |
|---|---|---|
| | muP*comX*-V3 | GAACggtctctATAGaactgtttaatctgttagtcagaggttgtaaataaaacagccagttaagatgTgacatttatgtcGtgttcttaaagtctttttcgttttataataattttattataaaaggaggtcatcgtaatagATGGAgagaccTTC |
| | muP*comS2*-V1 | GAACggtctctATAGtttcaaagtaaaaggggaagtcactagtacaaaaaggatgattctctttagaaaaatattctttttccaagaggGgacactaatgtcgttgttgctttgaggaatctggggtagaatagactcatcaaatgaaaggaagtctttatcaaATGGAgagaccTTC |
| | muP*comS2*-V2 | GAACggtctctATAGtttcaaagtaaaaggggaagtcactagtacaaaaaggatgattctctttagaaaaatattctttttccaagaggtgacactaatgtcCttgttgctttgaggaatctggggtagaatagactcatcaaatgaaaggaagtctttatcaaATGGAgagaccTTC |
| | muP*comS2*-V3 | GAACggtctctATAGtttcaaagtaaaaggggaagtcactagtacaaaaaggatgattctctttagaaaaatattctttttccaagaggGgacactaatgtcCttgttgctttgaggaatctggggtagaatagactcatcaaatgaaaggaagtctttatcaaATGGAgagaccTTC |
| | 6715P*comS*-V | GAACggtctctATAGtaaagtttctacggagcttcattgtatagagaagagtgatttcttttaagagatcacctttttagtttttggaaaTAgtgacataaatgtcgttttgttttgagctgaatgggtgtagaataggtttataaaatgaaagggagtttaataATGGAgagaccTTC |
| | O1-P*comX*rep-U-R | GAAGGTCTCTCTATaaatgaagaaacaacagcaaataaggact |
| | muP*comX*-V4 | GAACggtctctATAGaactgtttaatctgttagtcagaggttgtaaataaaacagccagttaagaGgTgacatttatgtcGtgttcttaaagtctttttcgttttataataattttattataaaaggaggtcatcgtaatagATGGAgagaccTTC |
| | muP*comX*-V5 | GAACggtctctATAGaactgtttaatctgttagtcagaggttgtaaataaaacagccagttaagatgTgacatttatgtcGtTttcttaaagtctttttcgttttataataattttattataaaaggaggtcatcgtaatagATGGAgagaccTTC |
| | muP*comX*-V6 | GAACggtctctATAGaactgtttaatctgttagtcagaggttgtaaataaaacagccagttaagaGgTgacatttatgtcGtTttcttaaagtctttttcgttttataataattttattataaaaggaggtcatcgtaatagATGGAgagaccTTC |
| | 6715P*comS*-V2 | GAACggtctctATAGtaaagtttctacggagcttcattgtatagagaagagtgatttcttttaagagatcacctttttagtttttggaaaaggtgacataaatgtcACtttgttttgagctgaatgggtgtagaataggtttataaaatgaaagggagtttaataATGGAgagaccTTC |
| | 6715P*comS*-V3 | GAACggtctctATAGtaaagtttctacggagcttcattgtatagagaagagtgatttcttttaagagatcacctttttagtttttggaaaaggtgacatTGatgtcgttttgttttgagctgaatgggtgtagaataggtttataaaatgaaagggagtttaataATGGAgagaccTTC |
| | 6715P*comS*-Va | GAACggtctctATAGtaaagtttctacggagcttcattgtatagagaagagtgatttcttttaagagatcacctttttagtttttggaaatagtgacattgatgtcactatgttttgagctgaatgggtgtagaataggtttataaaatgaaagggagtttaataATGGAgagaccTTC |
| | 67P*comX*-Va | GAACggtctctATAGccttaaagtagtcaaggctaatctattgctgtccaatttcttttaaggtgacataaatgtcgtttccaaatttttcagagtcctattataatttaagaaaaaatttaaaggagaatttagATGGAgagaccTTC |
| | 67P247 | GAACggtctctATAGactccaatcaactatccgattggagtttcgttttgttttctgcagaaggtggacctcttgataaagcgtgtttcttatgaagggggacattaatgtcagggtgacatcaatgtcactagtctttttcattgacttcttctataatgtaaccatagtgataaagaaaacatatcactaatttatttaaattgggtatagtggaaaggagattgttATGGAgagaccTTC |
| | 67P255 | GAACggtctctATAGttgaagctcctagaggttgttcgtccgataaaaggaaagggggtgacattgatgtcgttcctcttttggataatatcttttataataaggccataaaaaaggttatcttataccgaattaattggaaggaaaataATGGAgagaccTTC |

#### Supplementary Table 3. List of plasmids

| **Plasmids** | **Characteristics** | **Source** |
|---|---|---|
| pRW17 | Golden Gate assembly-compatible plasmid | (Li, et al., 2021) |
| **Plasmids for *S. sobrinus* (NIDR 6715-7)** | | |
| p67P*comS*-mNG | Spcr; Reporter for *S. sobrinus* P*comS* | This study |
| p67R'-P*comS*-mNG | Spcr; Reporter for *S. sobrinus* P*comS* with PcomR1 & part of comR1 | This study |
| p67P*comX*-mNG | Spcr; Reporter for *S. sobrinus* P*comX* | This study |
| p67PR1-mNG | Spcr; Reporter for *S. sobrinus* PcomR1 | This study |
| p67PR2-mNG | Spcr; Reporter for *S. sobrinus* PcomR2 | This study |
| p67P231-mNG | Spcr; Reporter for *S. sobrinus* P231, , unannotated between DLJ51_00230 & _00235 | This study |
| p67P4555-mNG | Spcr; Reporter for *S. sobrinus* P_DLJ51_04555 | This study |
| p67P4575a-mNG | Spcr; Reporter for *S. sobrinus* P_DLJ51_04575 | This study |
| p67P4575b-mNG | Spcr; Reporter for *S. sobrinus* P_DLJ51_04575, longer seqeunce taken | This study |
| p67P*cinA*-mNG | Spcr; Reporter for *S. sobrinus* P*cinA* | This study |
| p67P*comS*-V1 | Spcr; Reporter for *S. sobrinus* P*comS*-V1 | This study |
| p67P*comS*-V2 | Spcr; Reporter for *S. sobrinus* P*comS*-V2 | This study |
| p67P*comS*-V3 | Spcr; Reporter for *S. sobrinus* P*comS*-V3 | This study |
| p67P*comS*-Va | Spcr; Reporter for *S. sobrinus* P*comS*-Va | This study |
| p67P*comX*-Va | Spcr; Reporter for *S. sobrinus* P*comX*-Va | This study |
| p67P247 | Spcr; Reporter for *S. sobrinus* P247, unannotated between DLJ51_00245 & DLJ51_00250 | This study |
| p67P255 | Spcr; Reporter for *S. sobrinus* P255 | This study |
| **Plasmids for *S. downei*** | | |
| pDowP*comS1*-mNG | Spcr; Reporter for *S. downei* P*comS1* | This study |
| pDowP*comS2*-mNG | Spcr; Reporter for *S. downei* P*comS2* | This study |
| pDowR1'-P*comS1*-mNG | Spcr; Reporter for *S. downei* P*comS1* with PcomR1 & part of comR1 | This study |
| pDowR2'-P*comS2*-mNG | Spcr; Reporter for *S. downei* P*comS2* with PcomR2 & part of comR2 | This study |
| pDowP*comX*-mNG | Spcr; Reporter for *S. downei* P*comX* | This study |
| pDowPR1-mNG | Spcr; Reporter for *S. downei* PcomR1 | This study |
| pDowPR2-mNG | Spcr; Reporter for *S. downei* PcomR2 | This study |
| pDowP417-mNG | Spcr; Reporter for *S. downei* P417, unannotated between *comS1* & NCTC11391_00418 | This study |
| pDowP171-mNG | Spcr; Reporter for *S. downei* P_NCTC11391_00171 | This study |
| pDowPfasA2-mNG | Spcr; Reporter for *S. downei* PfasA_2 | This study |
| pDowPubaA-mNG | Spcr; Reporter for *S. downei* PubaA | This study |
| pDowPyfaY-mNG | Spcr; Reporter for *S. downei* PyfaY | This study |

| | | |
|---|---|---|
| **Plasmids for *S. criceti*** | | |
| pCriP*comS*-mNG | Spcr; Reporter for *S. criceti* P*comS* | This study |
| pCriR'-P*comS*-mNG | Spcr; Reporter for *S. criceti* P*comS* with PcomR & part of comR | This study |
| pCriP*comX*-mNG | Spcr; Reporter for *S. criceti* P*comX* | This study |
| pCriPsakP2-mNG | Spcr; Reporter for *S. criceti* PsakP2 | This study |
| pCriP*cinA*-mNG | Spcr; Reporter for *S. criceti* P*cinA* | This study |
| **Plasmids for *S. mutans*** | | |
| pUP*comS1*-mNG | Spcr; Reporter for *S. mutans* P*comS1* with mNeonGreen | This study |
| pUP*comS1*-mSC | Spcr; Reporter for *S. mutans* P*comS1* with mScarlet-I | This study |
| pUP*comS1*-mTB2 | Spcr; Reporter for *S. mutans* P*comS1* with mTagBFP2 | This study |
| pUP*comS1*-sfGFP | Spcr; Reporter for *S. mutans* P*comS1* with sfGFP | This study |
| pUP*comS1*-mTFP1 | Spcr; Reporter for *S. mutans* P*comS1* with mTFP1 | This study |
| pUP*comS1*-mTq2 | Spcr; Reporter for *S. mutans* P*comS1* with mTurquoise2 | This study |
| pUP*comS2*-mNG | Spcr; Reporter for *S. mutans* P*comS2* | This study |
| pUR1'-P*comS1*-mNG | Spcr; Reporter for *S. mutans* P*comS1* with PcomR1 & part of comR1 | This study |
| pUR2'-P*comS2*-mNG | Spcr; Reporter for *S. mutans* P*comS2* with PcomR2 & part of comR2 | This study |
| pUP*comX*-mNG | Spcr; Reporter for *S. mutans* P*comX* | This study |
| pUP*nlmA*-mNG | Spcr; Reporter for *S. mutans* P*nlmA* | This study |
| pUP370-mNG | Spcr; Reporter for *S. mutans* P_SMU.370 | This study |
| pUP*cinA*-mNG | Spcr; Reporter for *S. mutans* P*cinA* | This study |
| pUPcomE-mNG | Spcr; Reporter for *S. mutans* PcomE | This study |
| pUP*comX*-V1 | Spcr; Reporter for *S. mutans* P*comX*-V1 | This study |
| pUP*comX*-V2 | Spcr; Reporter for *S. mutans* P*comX*-V2 | This study |
| pUP*comX*-V3 | Spcr; Reporter for *S. mutans* P*comX*-V3 | This study |
| pUP*comS2*-V1 | Spcr; Reporter for *S. mutans* P*comS2*-V1 | This study |
| pUP*comS2*-V2 | Spcr; Reporter for *S. mutans* P*comS2*-V2 | This study |
| pUP*comS2*-V3 | Spcr; Reporter for *S. mutans* P*comS2*-V3 | This study |
| pUP*comX*-V4 | Spcr; Reporter for *S. mutans* P*comX*-V4 | This study |
| pUP*comX*-V5 | Spcr; Reporter for *S. mutans* P*comX*-V5 | This study |
| pUP*comX*-V6 | Spcr; Reporter for *S. mutans* P*comX*-V6 | This study |
| **Plasmids for *S. ratti*** | | |
| pRatP*comS1*-mNG | Spcr; Reporter for *S. ratti* P*comS1* | This study |
| pRatP*comS2*-mNG | Spcr; Reporter for *S. ratti* P*comS2* | This study |
| pRatR1'-P*comS1*-mNG | Spcr; Reporter for *S. ratti* P*comS1* with PcomR1 & part of comR1 | This study |

| | | |
|---|---|---|
| pRatR2'-P*comS2*-mNG | Spcr; Reporter for *S. ratti* P*comS2* with PcomR2 & part of comR2 | This study |
| pRatP*comX*-mNG | Spcr; Reporter for *S. ratti* P*comX* | This study |
| **Plasmids for *S. ferus*** | | |
| pFerP*comS1*-mNG | Spcr; Reporter for *S. ferus* P*comS1* | This study |
| pFeP*comS2*-mNG | Spcr; Reporter for *S. ferus* P*comS2* | This study |
| pFerR1'-P*comS1*-mNG | Spcr; Reporter for *S. ferus* P*comS1* with PcomR1 & part of comR1 | This study |
| pFerR2'-P*comS2*-mNG | Spcr; Reporter for *S. ferus* P*comS2* with PcomR2 & part of comR2 | This study |
| pFerP*comX*-mNG | Spcr; Reporter for *S. ferus* P*comX* | This study |

**Supplementary Table 4. List of strains**

| **Strains** | **Source** |
|---|---|
| *S. sobrinus* NIDR 6715-7 | ARS Culture Collection |
| *S. mutans* UA159 | ATCC |
| *S. ratti* FA1 | DSMZ |
| *S. ferus* 8S1 | DSMZ |
| *S. macacae* 25-1 | DSMZ |
| *S. downei* MFe28 (NCTC 11391) | DSMZ |
| *S. criceti* HS-6 | DSMZ |
| **Strains with deletion only** | |
| *S. sobrinus* ΔcomR1::CmR | (Li, et al., 2021) |
| *S. sobrinus* ΔcomR2::CmR | (Li, et al., 2021) |
| *S. sobrinus* Δ*comX*::CmR | This study |
| *S. downei* ΔcomRS1::CmR | This study |
| *S. downei* ΔcomRS2::CmR | This study |
| *S. downei* Δ*comX*::CmR | This study |
| *S. criceti* ΔcomRS::CmR | This study |
| *S. criceti* Δ*comX*::CmR | This study |
| *S. mutans* ΔcomRS1::CmR | This study |
| *S. mutans* ΔcomRS2::CmR | This study |
| *S. mutans* Δ*comX*::CmR | This study |
| *S. mutans* ΔcomDE::CmR | This study |
| *S. mutans* ΔSMU.370::CmR | This study |
| *S. mutans* ΔcomDE::CmR ΔSMU.370::KanR | This study |
| *S. ratti* ΔcomRS1::CmR | This study |
| *S. ratti* ΔcomRS2::CmR | This study |
| *S. ratti* Δ*comX*::CmR | This study |
| *S. ferus* ΔcomRS1::CmR | This study |
| *S. ferus* ΔcomRS2::CmR | This study |
| *S. ferus* Δ*comX*::CmR | This study |
| **Strains with reporter only** | |
| *S. sobrinus* p67P*comS*-mNG | This study |
| *S. sobrinus* p67R'-P*comS*-mNG | This study |
| *S. sobrinus* p67P*comX*-mNG | This study |
| *S. sobrinus* p67PR1-mNG | This study |
| *S. sobrinus* p67PR2-mNG | This study |

| | |
|---|---|
| *S. sobrinus* p67P231-mNG | This study |
| *S. sobrinus* p67P4555-mNG | This study |
| *S. sobrinus* p67P4575a-mNG | This study |
| *S. sobrinus* p67P4575b-mNG | This study |
| *S. sobrinus* p67P*cinA*-mNG | This study |
| *S. sobrinus* p67P*comS*-V1 | This study |
| *S. sobrinus* p67P*comS*-V2 | This study |
| *S. sobrinus* p67P*comS*-V3 | This study |
| *S. sobrinus* p67P*comS*-Va | This study |
| *S. sobrinus* p67P*comX*-Va | This study |
| *S. sobrinus* p67P247 | This study |
| *S. sobrinus* p67P255 | This study |
| *S. downei* pDowP*comS1*-mNG | This study |
| *S. downei* pDowP*comS2*-mNG | This study |
| *S. downei* pDowR1'-P*comS1*-mNG | This study |
| *S. downei* pDowR2'-P*comS2*-mNG | This study |
| *S. downei* pDowP*comX*-mNG | This study |
| *S. downei* pDowPR1-mNG | This study |
| *S. downei* pDowPR2-mNG | This study |
| *S. downei* pDowP417-mNG | This study |
| *S. downei* pDowP171-mNG | This study |
| *S. downei* pDowPfasA2-mNG | This study |
| *S. downei* pDowPubaA-mNG | This study |
| *S. downei* pDowPyfaY-mNG | This study |
| *S. criceti* pCriP*comS*-mNG | This study |
| *S. criceti* pCriR'-P*comS*-mNG | This study |
| *S. criceti* pCriP*comX*-mNG | This study |
| *S. criceti* pCriPsakP2-mNG | This study |
| *S. criceti* pCriP*cinA*-mNG | This study |
| *S. mutans* pUP*comS1*-mNG | This study |
| *S. mutans* pUP*comS1*-mSC | This study |
| *S. mutans* pUP*comS1*-mTB2 | This study |
| *S. mutans* pUP*comS1*-sfGFP | This study |
| *S. mutans* pUP*comS1*-mTFP1 | This study |
| *S. mutans* pUP*comS1*-mTq2 | This study |
| *S. mutans* pUP*comS1*-mNG | This study |
| *S. mutans* pUR1'-P*comS2*-mNG | This study |
| *S. mutans* pUR2'-P*comS2*-mNG | This study |
| *S. mutans* pUP*comX*-mNG | This study |

| | |
|---|---|
| *S. mutans* pUP*nlmA*-mNG | This study |
| *S. mutans* pUP370-mNG | This study |
| *S. mutans* pUP*cinA*-mNG | This study |
| *S. mutans* pUPcomE-mNG | This study |
| *S. mutans* pUP*comX*-V1 | This study |
| *S. mutans* pUP*comX*-V2 | This study |
| *S. mutans* pUP*comX*-V3 | This study |
| *S. mutans* pUP*comS2*-V1 | This study |
| *S. mutans* pUP*comS2*-V2 | This study |
| *S. mutans* pUP*comS2*-V3 | This study |
| *S. mutans* pUP*comX*-V4 | This study |
| *S. mutans* pUP*comX*-V5 | This study |
| *S. mutans* pUP*comX*-V6 | This study |
| *S. ratti* pRatP*comS1*-mNG | This study |
| *S. ratti* pRatP*comS2*-mNG | This study |
| *S. ratti* pRatR1'-P*comS1*-mNG | This study |
| *S. ratti* pRatR2'-P*comS2*-mNG | This study |
| *S. ratti* pRatP*comX*-mNG | This study |
| *S. ferus* pFerP*comS1*-mNG | This study |
| *S. ferus* pFeP*comS2*-mNG | This study |
| *S. ferus* pFeR1'-P*comS1*-mNG | This study |
| *S. ferus* pFeR2'-P*comS2*-mNG | This study |
| *S. ferus* pFeP*comX*-mNG | This study |
| **Reporter + deletions** | |
| *S. sobrinus* ΔcomR1::CmR p67P*comS*-mNG | This study |
| *S. sobrinus* ΔcomR2::CmR p67P*comS*-mNG | This study |
| *S. sobrinus* ΔcomR1::CmR p67P231-mNG | This study |
| *S. sobrinus* ΔcomR2::CmR p67P231-mNG | This study |
| *S. sobrinus* ΔcomR1::CmR p67P*comS*-V1 | This study |
| *S. sobrinus* ΔcomR2::CmR p67P*comS*-V1 | This study |
| *S. sobrinus* ΔcomR1::CmR p67P*comS*-V2 | This study |
| *S. sobrinus* ΔcomR2::CmR p67P*comS*-V2 | This study |
| *S. sobrinus* ΔcomR1::CmR p67P*comS*-V3 | This study |
| *S. sobrinus* ΔcomR2::CmR p67P*comS*-V3 | This study |
| *S. sobrinus* ΔcomR1::CmR p67P*comS*-Va | This study |
| *S. sobrinus* ΔcomR2::CmR p67P*comS*-Va | This study |
| *S. sobrinus* ΔcomR1::CmR p67P*comX*-Va | This study |
| *S. sobrinus* ΔcomR2::CmR p67P*comX*-Va | This study |

| | |
|---|---|
| *S. sobrinus* ΔcomR1::CmR p67P247 | This study |
| *S. sobrinus* ΔcomR2::CmR p67P247 | This study |
| *S. sobrinus* ΔcomR1::CmR p67P255 | This study |
| *S. sobrinus* ΔcomR2::CmR p67P255 | This study |
| *S. mutans* ΔcomRS1::CmR pUP*comS1*-mNG | This study |
| *S. mutans* ΔcomRS2::CmR pUP*comS1*-mNG | This study |
| *S. mutans* Δ*comX*::CmR pUP*comS1*-mNG | This study |
| *S. mutans* ΔcomRS1::CmR ΔcomRS2::KanR pUP*comS1*-mNG | This study |
| *S. mutans* ΔcomRS1::CmR pUP*comS2*-mNG | This study |
| *S. mutans* ΔcomRS2::CmR pUP*comS2*-mNG | This study |
| *S. mutans* Δ*comX*::CmR pUP*comS2*-mNG | This study |
| *S. downei* ΔcomRS1::CmR pDowP*comS1*-mNG | This study |
| *S. downei* ΔcomRS2::CmR pDowP*comS1*-mNG | This study |
| *S. downei* ΔcomRS1::CmR pDowP*comS2*-mNG | This study |
| *S. downei* ΔcomRS2::CmR pDowP*comS2*-mNG | This study |
| *S. downei* ΔcomX::CmR pDowP*comS1*-mNG | This study |
| *S. downei* ΔcomX::CmR pDowP*comS2*-mNG | This study |

#### Supplementary Table 5. RODEO analysis of bacteriocin clusters predicted in the ComRS networks [1]

| Promoter | Predicted RBS strength | Encoding | Peptide Sequence |
|---|---|---|---|
| ***S. sobrinus* cluster A** | | | |
| | | Phosphate transferase, PlsX | |
| | | DLJ51_00230, Acyl carrier protein | |
| 2x ComR box | | | |
| | 21015 | Pep1 | MLKKKKGSEALVGQELAKV* |
| | 20913 | Pep2 | MNYVELTTDELAAIEGGNILHSIGNAFSSAYHGLVDGWNKH* |
| | 20150 | Pep3 | MDTTVEYSTLTVEELSELIGGNVFYDAAYQVGSFARGVWDAF * |
| | 23393 | Pep4 | METKTFEKYETVNVEELAELVGGNTAYNAGKVVGKVGQIAA VIAAFF** |
| | | DLJ51_00240, Bacitracin ATP Transporter | |
| | | DLJ51_00245, Permease | |
| | 1891 | Pep5 | VMQELQSTIRLEFRFVFCRRWTS** |
| 2x ComR box | | | |
| | 4088 | Pep6 | MSKENKNPQTLTDQDLAKVQGGNWINVFKSVVDIARRGVIVY * |
| | 979 | Pep7 | MFLNLLLTSLVGVLLFIRYSLVDRLLVLKSVSS** |
| | 3035 | Pep8 | LFIRYSLVDRLLVLKSVSS** |
| | 452 | Pep9 | MMVLIIFSERIVVSSLASGFIMKKMGSVIAFLLVAIMAVLLTLK FFSGQVSHSAPLVLVLIALIVLGKKALANIKS* |
| | 1846 | Pep10 | MLKLLEVVRPIKGKG* |
| | 1717 | Pep11 | MSFLFWIISFIIRP* |
| ComR box | | DLJ51_00255, Lipase | |
| | 24190 | Pep12 | MKTKELLFSELNEEDLAAIRGGSLWDFKGIIIGDPWWYPRIGIP EHPILLDK* |
| | | ComR | |
| ComR box | | ComS | MNLKKIIELAITLVALMCTIAR |
| | | Transporter ComA | |
| ***S. downei* cluster A** | | | |
| | | recO, plsX, acp | |
| | | ComR1 | |
| ComR box | | | |
| | 243 | *comS1* | MKKLFLILASLATMVLFTF* |
| | 69 | Pep1 | LIGSPYPLQGK** |
| | 90 | Pep2 | LPYLLRKAKLSLFPTSKGLPDLLSCLRLLKKGVTWMSGDIHVAI FLSRSIL* |
| 2x ComR box | | | |
| | 10449 | Pep3 | MERIIWGDEVYRFPPKFKRKLGF* |
| | 13530 | Pep4 | MAKKVKAKAGKRNTVNLGIVSYTYTRGRHSI* |
| | 15325 | Pep5 | MAKKVKAKASKRNTVNLGIVSYTYTRGRHSI* |
| | 22584 | Pep6 | MTKKVKAKSNKRNTVNLGIVSYTYTRGRHSI* |
| | 5521 | Pep7 | MAKKVKAKSNKRNTVNLGIVSYTYTRGRHSI* |
| | 11141 | Pep8 | MAKKVKAKASKRNTVNLGIVSYTYTRGRHSI* |
| | 9475 | Pep9 | MAKKVKAKASKRNTVNLGIVSYTYTRGHHSI* |
| | 30364 | Pep10 | MQKSNSGFKMLTDKQLSKISGGAGYVMPSDWPWTKFKWPRG VIPWK* |
| | 1372 | Pep11 | MSCRVIGPGPNSSGREGLFHGNEK* |
| | | *NCTC11391_00419*, amino acid transporter | |
| | | *NCTC11391_00420*, metal-dependent membrane protease | |
| | | *NCTC11391_00421*, membrane protein | |
| | | MetN_1, multidrug ABC transporter ATPase | |
| | | *NCTC11391_00423*, sugar ABC transporter permease | |
| | | LmrA, multidrug ABC transporter ATPase and permease | |
| | 30364 | Pep12 | MQKSNSGFKMLTDKQLSKISGGAGYVMPSDWPWTKFKWPRG VIPWK* |
| | 1372 | Pep13 | MSCRVIGPGPNSSGREGLFHGNEK* |
| | | lipase | |
| | 6443 | Pep14 | VLGEKDLSKEQVWS* |

| | | | |
|---|---|---|---|
| | 172 | Pep15 | MARSSKGVSLLWDVQLISVIPI*** |
| | | comR2 | |
| 2x ComR box | | *comS2* | |
| | 2368 | Pep16 | MKLRKVLEILTVIAMLTVLFR* |
| | 1485 | Pep17 | LKKYSYVYQGLKGKIMKK* |
| ***S. sobrinus* B** | | | |
| cin-box | | *DLJ51_4555, DNA-binding response regulator* | |
| | | *DLJ51_4560, Histidine kinase* | |
| | | *DLJ51_4565, HlyD family secretion protein* | |
| | | *DLJ51_4570, peptide cleavage/export ABC transporter* | |
| | | *DLJ51_4575, ComC/BlpC family peptide pheromone/bacteriocin* | MMKTQAIEKFDVMDTEALSTVEGGRKWGNCEIAVAGGAVLG GAITGGNPLGAIGGAMMAAAEYC* |
| | | *DLJ51_4580, hypothetical protein* | MNVQLLDKFEAIEKDYLSTVKGGNWRCYVGTAGGAIFGAVG GPWGAIAGGTVANAYFCAEKVS* |
| | | *DLJ51_4585, hypothetical protein* | MDNLKHYTIIGIRILAVVVAISIGLKVLIYSTELWRKIFVIIGVPL FLYAITTVRRRKCTK* |
| | | *DLJ51_4590, IS256 family transposase* | |
| | | *DLJ51_4595, bacteriocin-type signal sequence* | MKFNFTNYENLSTKELSFIQGGDWRNELLNFLIETSMPIGWNW HQKK* |
| | | *DLJ51_4600, hypothetical protein* | |
| | | *DLJ51_4605, thiol reductase thioredoxin* | |
| | | *, IS200/IS605 family transposase* | |
| | | *DLJ51_4615, psuedogene* | |
| | | *DLJ51_4620, frameshifted immunity protein* | |
| | | *DLJ51_4625, hypothetical protein* | |
| | | *DLJ51_4630, bacteriocin immunity protein* | |
| | | *DLJ51_4635, bacteriocin immunity protein* | |
| | | *DLJ51_4640, hypothetical protein* | |
| | | *DLJ51_4645, efflux transporter periplasmic adaptor subunit* | |
| | | *DLJ51_4650, ABC transporter ATP-binding protein* | |
| | | *DLJ51_4655, hypothetical protein* | |
| | | *DLJ51_4660, PTS sugar transporter subunit IIC* | |
| | | *DLJ51_4665, alpha/beta hydrolase* | |
| | | *DLJ51_4670, short-chain dehydrogenase* | |
| | | *DLJ51_4675, nuclear transport factor 2 family protein* | |
| | | *DLJ51_4680, transcriptional regulator* | |
| ***S. downei* B** | | | |
| cin-box | | *fasA_2, NCTC11391_01284, FasA_2, response regulator* | |
| | | *dcuS/comD, DcuS/ComD, histidine kinase* | |
| | | *lcnD CDS , LcnD, BlpC ABC transporter* | |
| | | *lagD_3 CDS , LagD_3, competence factor transporting ATP-binding protein/permease ComA* | |
| | | *ubaA CDS, UbaA, bacteriocin ubericin-A* | MKTQAIERFEVMNSEVLSTVEGGRTIYYGNGVYCNKTCWVD WGQTINSIATNSAMNWATQGNAGWHSGGVA* |
| | | *NCTC11391_01289, putative bacteriocin immunity protein* | |
| | | *NCTC11391_01290, Pep2* | MNFYKFVPQIDARNEGGRWFRAAAENSKNDL* |
| | | *NCTC11391_01291, Bacteriocin class II, pep3* | MKVQAIERFDVMDTEALSTVEGGQASLGCVLGTAGMAGAGF VFGGGPAGAAVLGGATALRLCR* |
| | | *NCTC11391_01292, pep4* | MKNKVLNDLKSLDNITLENIYGGKINPVCAVAVAGSIYTATAF ASGGAGVPLVIGAGAAAAAGFCG* |
| | | *NCTC11391_01293, ?* | |
| | | *NCTC11391_01294, putative bacteriocin* | |
| | | *NCTC11391_01295, pep5* | MKTQAIEKFYVVNDKQLETINGGVVGTIIAAGGAVMAGFQFA HWLGQMQAESDYNRTHR* |
| | | *NCTC11391_01296, pep6* | MINIDKFDQLNANDLSNCIGGYCGRYGLIAYGLGGALTGSFYC TGYENHYREMQRNYGY* |
| | | *NCTC11391_01297, pep7* | |
| | | *NCTC11391_01298, Xre-like regulator?* | |
| | | *NCTC11391_01299, ?* | |
| | | *NCTC11391_01300, ?* | |
| | | *NCTC11391_01301, membrane protein* | |
| | | *NCTC11391_01302, ABC transporter substrate-binding protein* | |
| | | *macB_4 CDS , macB_4, lipoprotein-releasing system ATP-binding protein* | |

| | | | |
|---|---|---|---|
| | | *macB_5 CDS , macB_5, ABC transporter permease* | |
| | | *chbC, ChbC, PTS cellobiose-specific IIC component* | |
| | | *NCTC11391_01306, cell surface hydrolase* | |
| | | *ydfG_1, YdfG_1, utative oxidoreductase* | |
| | | *NCTC11391_01308, SnoaL-like domain* | |
| | | *NCTC11391_01309, transcriptional regulator* | |
| ***S. downei* C** | | | |
| Double ComR box | | *NCTC11391_00171, thiazolylpeptide-type bacteriocin precursor* | MKDENKLLEELEIDILELDDVEGAPATAASSGSGGSSTCGSTSCSSCSSSCA* |
| | | *NCTC11391_00170, thiazolylpeptide-type bacteriocin precursor* | MKDENKLLEELEIDILELDDVEGVPATAATSGSGGSSTCGSTSCSSCSSSCA* |
| | | *NCTC11391_00169 , Uncharacterised protein* | |
| | | *NCTC11391_00168, bacteriocin biosynthesis docking scaffold, SagD family* | |
| | | *ssuB, ABC transporter ATP-binding protein* | |
| | | *NCTC11391_00166, ABC transporter* | |
| | | *NCTC11391_00165 , peptidase* | |
| | | *NCTC11391_00164, Lantibiotic biosynthesis protein* | |
| | | *NCTC11391_00163 , Uncharacterised protein* | |
| | | *NCTC11391_00162, Uncharacterised protein* | |
| | | *NCTC11391_00161, bacteriocin biosynthesis docking scaffold, SagD family* | |
| | | *NCTC11391_00160, thiopeptide-type bacteriocin biosynthesis domain* | |
| | | *sagB, streptolysin S biosynthesis protein* | |
| | | *NCTC11391_00158 , Uncharacterised protein* | |
| ***S. mutans* B** | | | |
| 2x ComR box | | putative ABC transporter, ATP-binding protein | |
| | | conserved hypothetical protein; inner membrane protein | |
| | | hypothetical protein | |
| | | hypothetical protein | |
| | | putative oxidoreductase | |
| | | hypothetical protein | |
| | | putative aminotransferase | |
| | 6418 | Pep1 | MEEKKQMSNVNLYQEIEVDELNGWGSYALGVAVGAGGSLAVAIT* |
| | | hypothetical protein | |
| | 1476 | Pep2 | LQILKQIKKLKRRRKKMEKELMFEEVTVEELNGINLNDFNNGVIAGAAVVAAGAGLAAVLT* |
| | 366860 | Pep3 | MEKELMFEEVTVEELNGINLNDFNNGVIAGAAVVAAGAGLAAVLT* |
| ComR box | 8214 | ComS2 (reverse strand) | MNLKKVLEILTTLVMLVMAVR*** |
| | | ComR2 (reverse strand) | |
| ***S. ratti* B** | | | |
| ComR box | | FY406_03900 (ABC transporter ATP-binding protein) | |
| | | FY406_03905 CDS (ABC transporter permease) | |
| | | FY406_03910 CDS (hypothetical protein) | |
| | | FY406_03915 CDS (class I SAM-dependent methyltransferase) | |
| | | FY406_03920 CDS (SDR family oxidoreductase) | |
| | | FY406_03925 CDS (hypothetical protein) | |
| | | FY406_03930 CDS (aminotransferase class III-fold pyridoxal phosphate-dependent enzyme) | |
| | | | VQKENDTELQIFEEVTVEELNTNFGKLIWKALLQPVIL* |
| | | | MEEKAFQYVEFEEISSEEFNGISVGQVAAFGAGVAVGALVIT* |
| | | | MKEHNETKMTAVQFEEINSEELNGDGWFIAGAVTGAGAVVGVAILT* |
| | | | MINSEPKNYFESILLLTAATILMIETKHLLVAAVMLVLTLLLSILGRKKVVSKSSRTNNKEKQ* |
| | | | MNNLDISTTEFQAETMFEELNSVELNGAARDFILGAAAGATIAGAIIT* |
| ***S. ferus* B** | | | |
| ComR box | | DQL21_RS08775 putative ABC transporter, ATP-binding protein | |
| | | DQL21_RS08770 inner membrane protein | |
| | 162411 | | MLSVIIMPGIWLAKWLF* |
| | 11794 | | MNGILDWFINHRIYIIC** |

| | | | |
|---|---|---|---|
| | | DQL21_RS08765 to SMU_372 | |
| | | DQL21_RS08760 to SMU_373 | |
| | | DQL21_RS08755 to SMU_374 CDS (putative oxidoreductase) | |
| | | DQL21_RS08750 to SMU_375 | |
| | | DQL21_RS08745 to SMU_376 CDS (putative aminotransferase) | |
| | 212003 | | MEKELITFEEVDSKEFNGWAEIGIAFGTGVVIGGAIVLT* |
| | 11477 | | VKDSKNSSLIKLQSVFLTISLLGTLAIILAESRLLVSLIIMVVIIILAFTNMLWLRQLDNKVNNR* |
| | 1418 | | MELQNNQVLVFEEVETAELNGNARDAVTGFGVGIGIVAAAAGIVALT* |
| | 146422 | | MELTNSFTKELVFEEVETAELNGNARDAIEGFGVGIGIVAAAAGIVALT* |
| | 55117 | | MELTDNLMNELTFEEVTTDELYGDGWFVGGVAGGVVVGGAIWAGVVLT* |
| ***S. macacae* B** | | | |
| ComR box | | STRMA_1436 CDS (ABC transporter, ATP-binding protein) | |
| | | STRMA_1435 CDS (putative membrane protein) | |
| | 97774, 312030 | | MMKGVLGWFFDHVQVIVY* |
| | | STRMA_1434 CDS (hypothetical protein)SMU_372 homolog | |
| | | STRMA_1433 CDS (methyltransferase domain protein) | |
| | | STRMA_1432 CDS (oxidoreductase, short chain dehydrogenase/reductase family protein) | |
| | | STRMA_1431 CDS (hypothetical protein) | |
| | | STRMA_1430 CDS (aminotransferase, class III) | |
| | directly following the above | | |
| | 1093 | | MKGGELMKLMQYFESLLLLNGAAIVLMQTNHATIAYLLLALSIVLSALGIKIARETK* |
| | 702036 | | MKLMQYFESLLLLNGAAIVLMQTNHATIAYLLLALSIVLSALGIKIARETK* |
| | 67461 | | MQKEFQDSQFVTFEEVDTTELNGAARDFILGAAAGATIAGAIIT* |
| | 45636 | | MQKTENNSINHNLFFEEMDSSELYGDGWFIAGAVTGAGVVVGAAILT* |
| | 33174 | | MKEQKAEMLFEEVDTTELNGAARDFILGAAAGATIAGAIIT* |
| | 39728 | | MQKTENNSINHNLFFEEMDSSELYGDGWFIAGAVTGAGVVVGAAILT* |
| | 2864 | | MHLTSFFSEIVDILLKVSHHLSKIDDLLAAIVA* |

**Supplementary Text. Reporter construction for ComRS network characterization**

As control by a ComRS system could be either direct or indirect, and endogenous activation could mix with effects of artificially-added inducers, we tried to use a fluorescence reporter with a short maturation time to best reflect the response time and distinguish these effects. Preliminary experiments showed that the P*comS1*$^{SMU}$ promoter has a quick and dramatic response to XIP1$^{SMU}$ addition. We thus chose this promoter as the test promoter to compare performance between different fluorescence reporters. Supplementary Figure 1A shows that among 6 candidate fluorescent proteins (mNeonGreen, mScarlet-1, sfGFP, mTurquoise2, mTFP1, and mTagBFP2), mNeonGreen and sfGFP showed the fastest response upon XIP1$^{SMU}$ supplementation (Supplementary Figure 1A). Compared with WT strain, the strain carrying mNeonGreen- or sfGFP-based P*comS1*$^{SMU}$ reporter plasmids also did not show noticeable growth defects, although the growth dynamics is moderately altered (Supplementary Figure 1B). Since we are mainly interested in the activation stages of the promoter activities, we chose to tolerate the potential changes to growth dynamics which likely mainly take effect after the initial activation stage. As mNeonGreen showed stronger signal magnitudes compared with sfGFP, which is beneficial for reflecting weaker promoter activities, we chose mNeonGreen as the reporter gene. Note that, as we used plasmid-encoded reporters, which are often multi-copy and with fluctuating copy number (Shao et al. 2021), it could only imprecisely approximate the true expression levels and dynamics of genes on bacterial chromosome. Additionally, it is also challenging to know how similar the degradation dynamics of the reporter protein is to the protein-of-interest. Nonetheless, the reporters are still valuable for showing whether a gene is induced by a XIP, the relative activation levels in different scenarios, and its activation time.

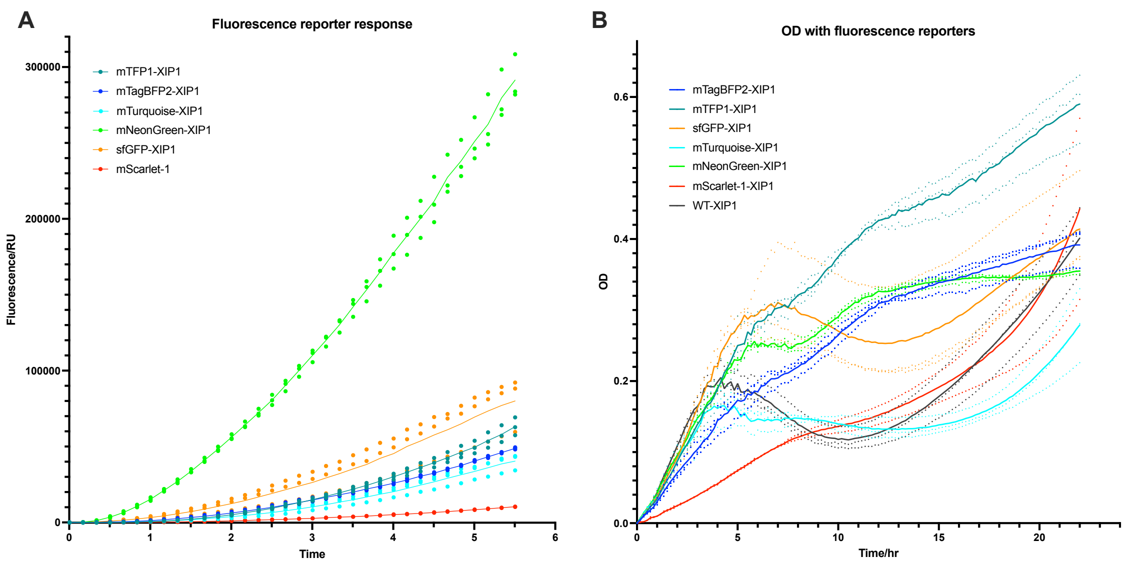

**Supplementary Figure 1. Fluorescent protein reporter selection by P*comS1*$^{SMU}$ response characteristics.** (**A**) Measured culture fluorescence after XIP1$^{SMU}$ is added for plasmid-encoded P*comS1*$^{SMU}$ reporters using difference fluorescent protein genes. Showing the first 5.5 hrs to highlight response time (protein maturation time). Time unit: hour. (**B**) OD of the corresponding cultures over 22 hrs for measuring growth effects from protein expression.

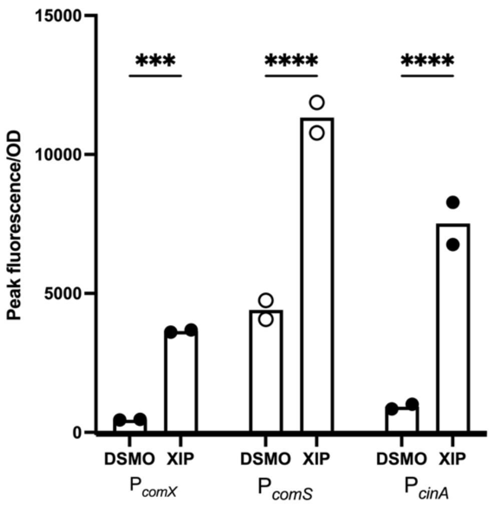

**Supplementary Figure 2. *S. criceti* promoter response to XIP.** Comparisons done with two-sample t tests. P-value notations: "ns", P > 0.05; "*", P < 0.05; "**", P < 0.01; "***", P < 0.001; "****", P < 0.0001. Same below.

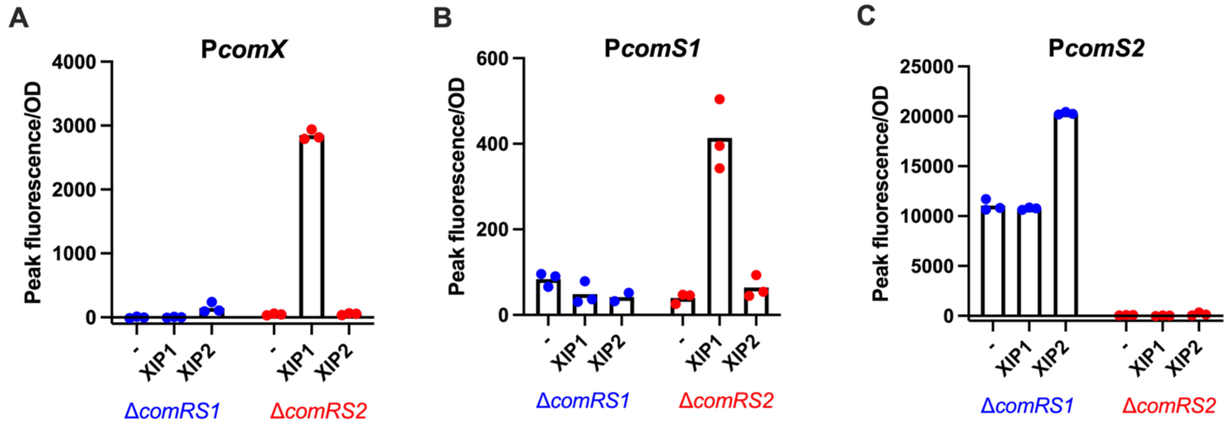

**Supplementary Figure 3. Response of (A) P*comX*, (B) P*comS1*, and (C) P*comS2* to XIPs in *S. downei* with either ComRS system deleted.**

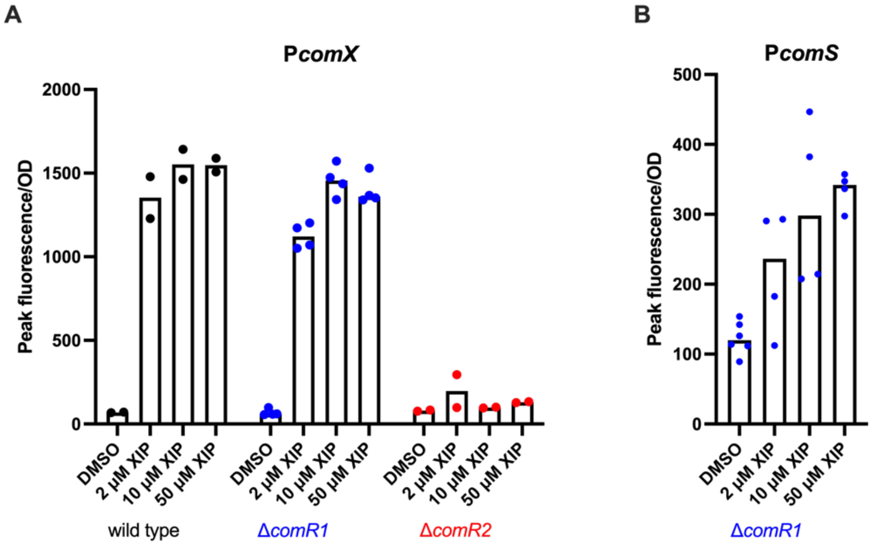

**Supplementary Figure 4. Concentration-dependent response of (A) P*comS* and (B) P*comX* to XIP in *S. sobrinus*.**

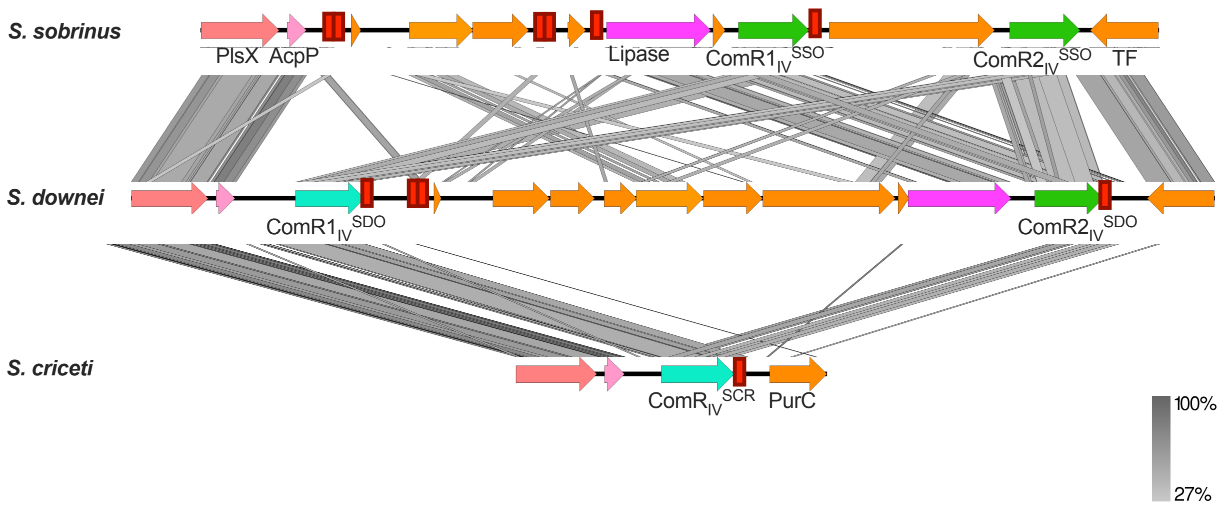

**Supplementary Figure 5. Alignment of *comRS* region between *S. downei* and *S. sobrinus*, *S. downei* and *S. criceti* based on encoded protein sequences.** The colors of ComR's correspond to sequence similarity (Figure 2). Figure made with EasyFig [2].

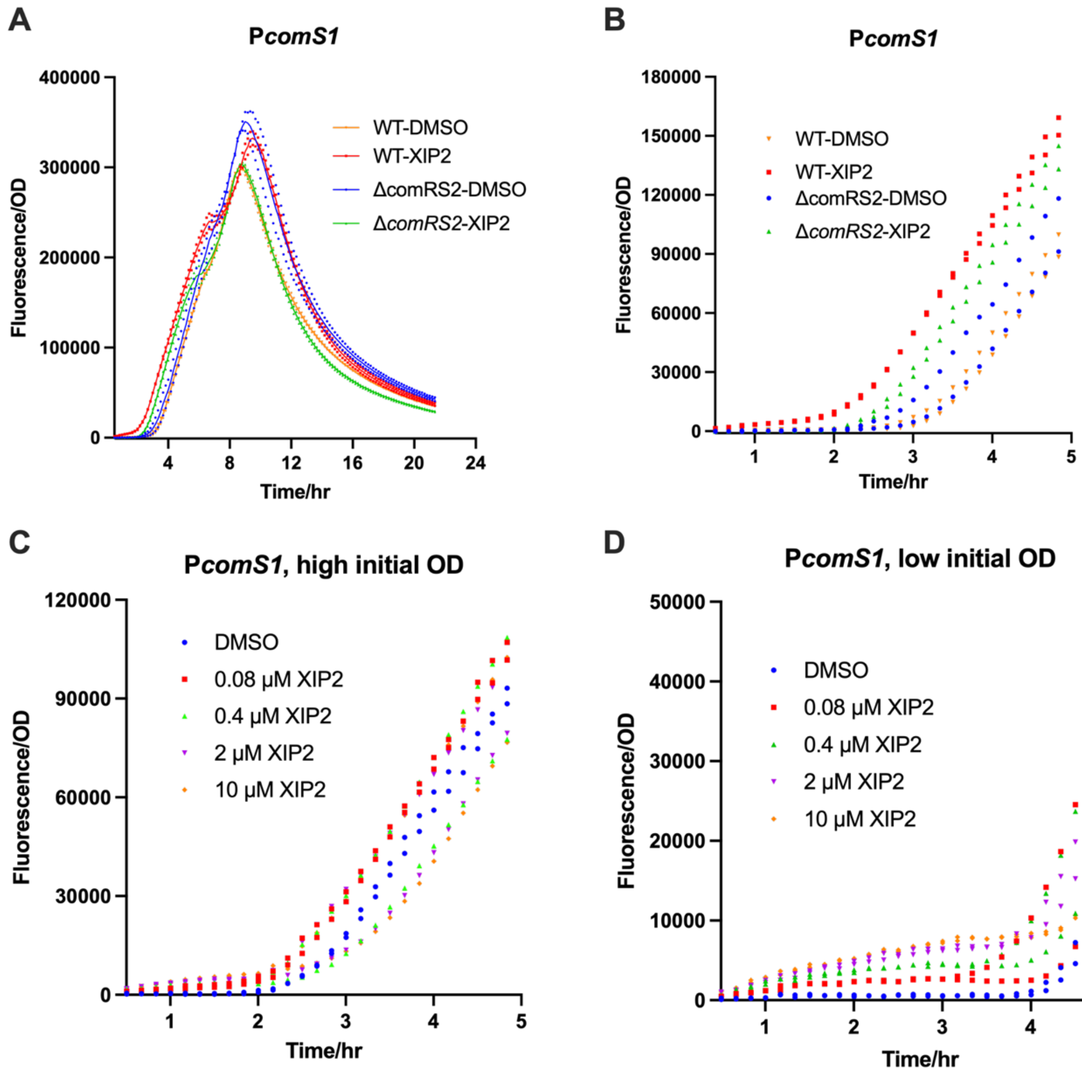

**Supplementary Figure 6. Closeup look at kickstart effects of XIP2 on P*comS1* through ComR2 in *S. mutans*. A&B** are based on the same dataset, showing that XIP2 can lead to early induction of P*comS1*, which depends on ComR2. **C&D.** The effect of XIP2 on P*comS1* is more pronounced when XIP2 is added at low OD (<0.15).

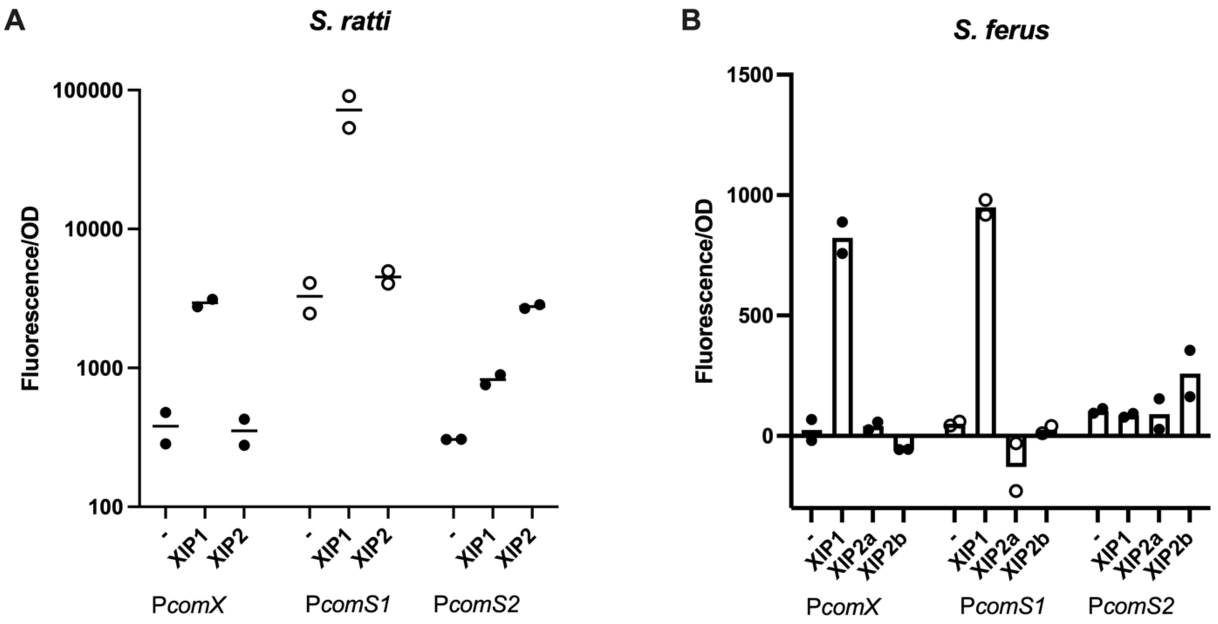

**Supplementary Figure 7. Response of P*comX*, P*comS1*, and P*comS2* to XIP in (A) *S. ratti* and (B) *S. ferus*.**

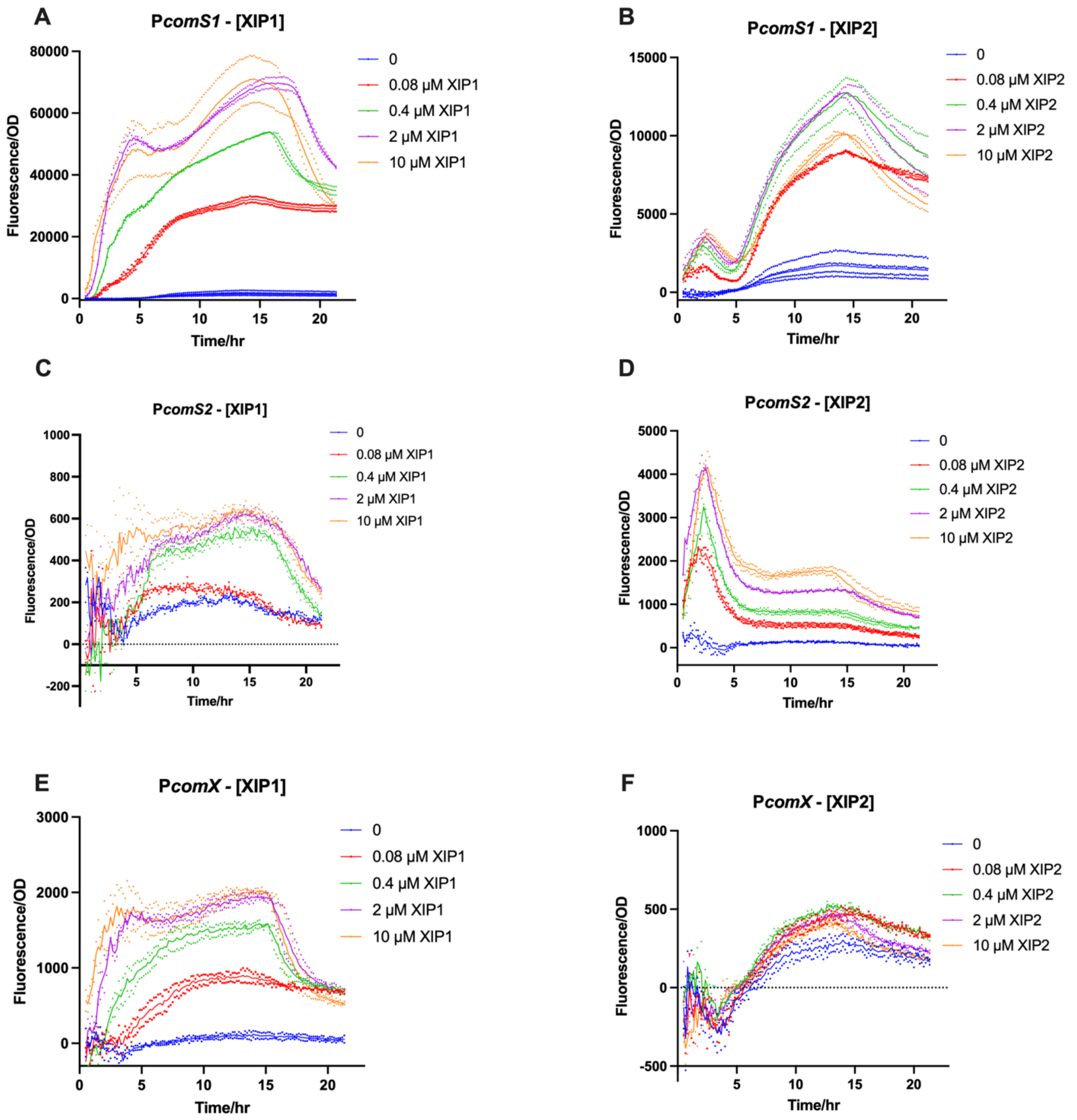

**Supplementary Figure 8. Concentration-dependent response of P*comX*, P*comS1*, and P*comS2* to XIPs in *S. ratti*.**

**A**

| Promoter | ComR box | Distance ComR to -10 | Controlling ComR-XIP |
|---|---|---|---|
| *PcomS1* | acgggacataaatgtcctgt | 20 | ComR1-XIP1 |
| *PcomX* | atgggacatttatgtcctgt | 20 | ComR1-XIP1 |
| *PcomS2* | aggtgacactaatgtcgttg | 20 | ComR2-XIP2 |
| P370-Box1 | tagtgacattaatgtcgtct | 20 | ComR2-XIP2 |
| P370-Box2 | gtctgacattgatgtcactt | 20 | ComR2-XIP2 |
|  | **** ***** |  |  |

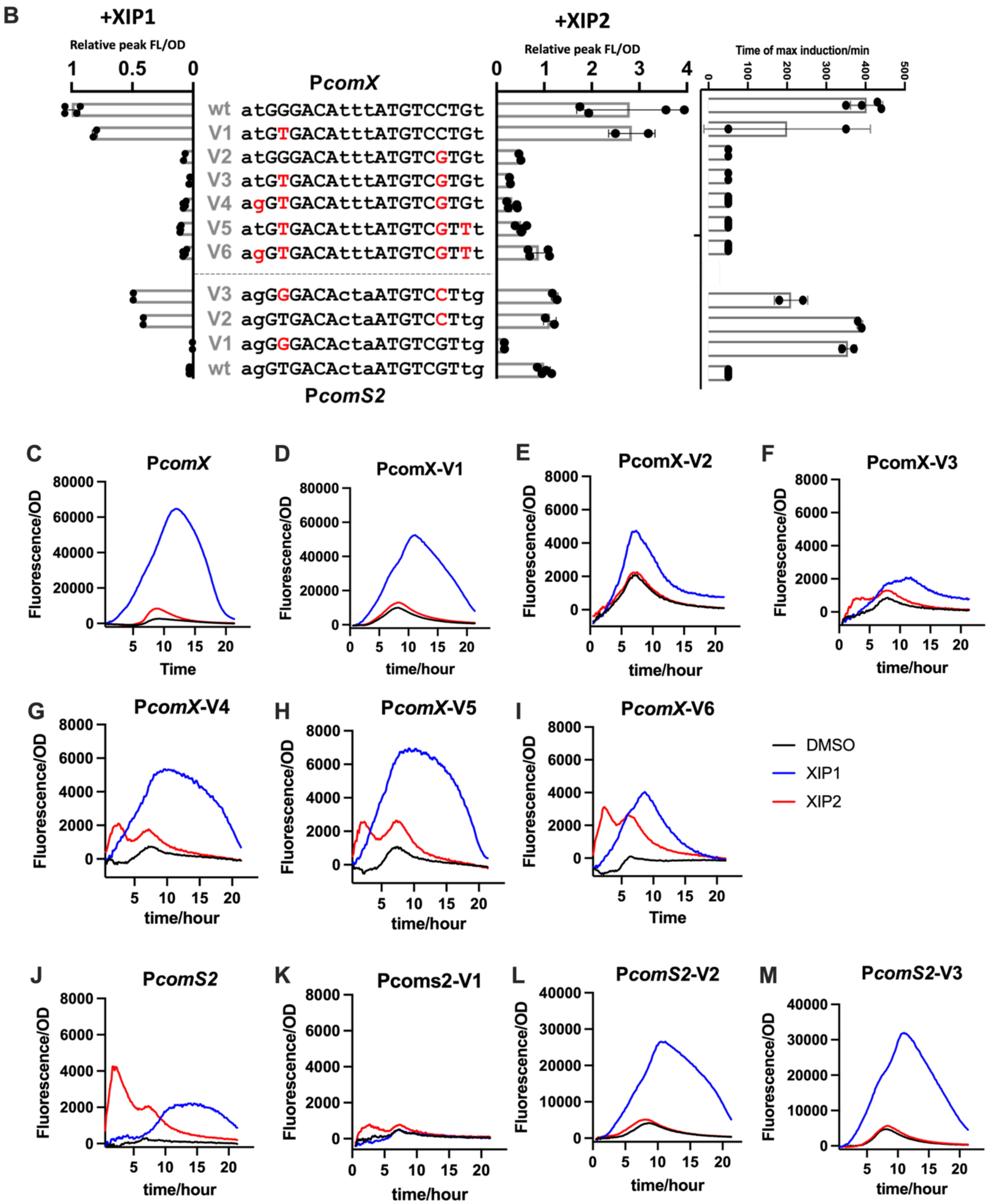

**Supplementary Figure 9. Extended analysis and data on effects of base pair changes in ComR boxes in *S. mutans.*** (**A**) ComR box feature analysis between ComRS1- and ComRS2-controlled promoter groups. P370 has double ComR boxes with 4-bp overlap. Red letters indicate consensus base pairs between P*comS1* and P*comX*. "*" indicates consensus among all boxes; "." on top indicates pyrimidine vs purine distinction between two groups. (**B**) Peak activity comparisons under XIP1 or XIP2 induction, and time (in min) of max induction rate after XIP2 addition, for different promoter variants. Note that as experiments are done in WT background, higher values don't necessarily indicate control by one system, since P*comX* has higher endogenous activity level than P*comS2* even when induced with XIP2. "P*comS2*-type" promoter should have high peak FL/OD with XIP2 induction as well as short response time. (**C-M**) Time course of reporter response to XIP1 and XIP2 for all promoter variants. Lines are shown as averages of duplicates (no noticeable variation seen between duplicates).

**A**

```
Promoter      ComR box                          Distance ComR box to -10    Controlling ComR-XIP
               .                   .
PcomS         aggtgacataaatgtcgttt                       20                       ComR1
P255          gggtgacattgatgtcgttc                       20                       ComR1
PcomX         tagtgacattgatgtcacta                       21                       ComR2
                *******  .*****   *

P231        atgcgacattaatgtcag-gtgacactaatgtctcta        21                       ComR1
P247        ggggacattaatgtcagggtgacatcaatgtcacta         21                  ComR1 & ComR2
            . * ************** ******   ****** ***
```

**B**

| Promoter | ComR box/sequence features | Controlling ComR |
|---|---|---|
| **P*comS*$_{IV}^{SSO}$** | aggtgacataaatgtcgttt | ComR1$_{IV}^{SSO}$-controlled |
| **P*comS*$_{IV}^{SSO}$-V1** | TAgtgacataaatgtcgttt | ComR1$_{IV}^{SSO}$-controlled |
| **P*comS*$_{IV}^{SSO}$-V2** | aggtgacataaatgtcACtt | Irresponsive to either ComR |
| **P*comS*$_{IV}^{SSO}$-Va** | *PcomX$^{SSO}$ box* + *PcomS$_{IV}^{SSO}$* surrounding | Irresponsive to either ComR |
| **P*comX*$^{SSO}$** | TAgtgacattgatgtcACta | ComR2$_{IV}^{SSO}$-controlled |
| **P*comX*$^{SSO}$-Va** | *PcomS$_{IV}^{SSO}$ box* + *PcomX$^{SSO}$* surrounding | Irresponsive to either ComR |

**Supplementary Figure 10. Analysis and summary data on effects of base pair changes in ComR boxes in *S. sobrinus*. (A)** Promoters P231 and P247 have double ComR boxes and are analyzed separately. Red shows consensus base pairs between P*comS* and P255, both controlled by ComR1. "*" indicate consensus among all three boxes; "." on *bottom* indicates pyrimidine/purine agreement between two groups; "." on *top* indicates pyrimidine vs purine distinction between two groups. **(B)** Summary of *S. sobrinus* promoter variant characteristics. Red: base pair changes. Capitalized: base pairs in P*comX* (ComR2-controlled) distinct from ComR1-controlled promoters.

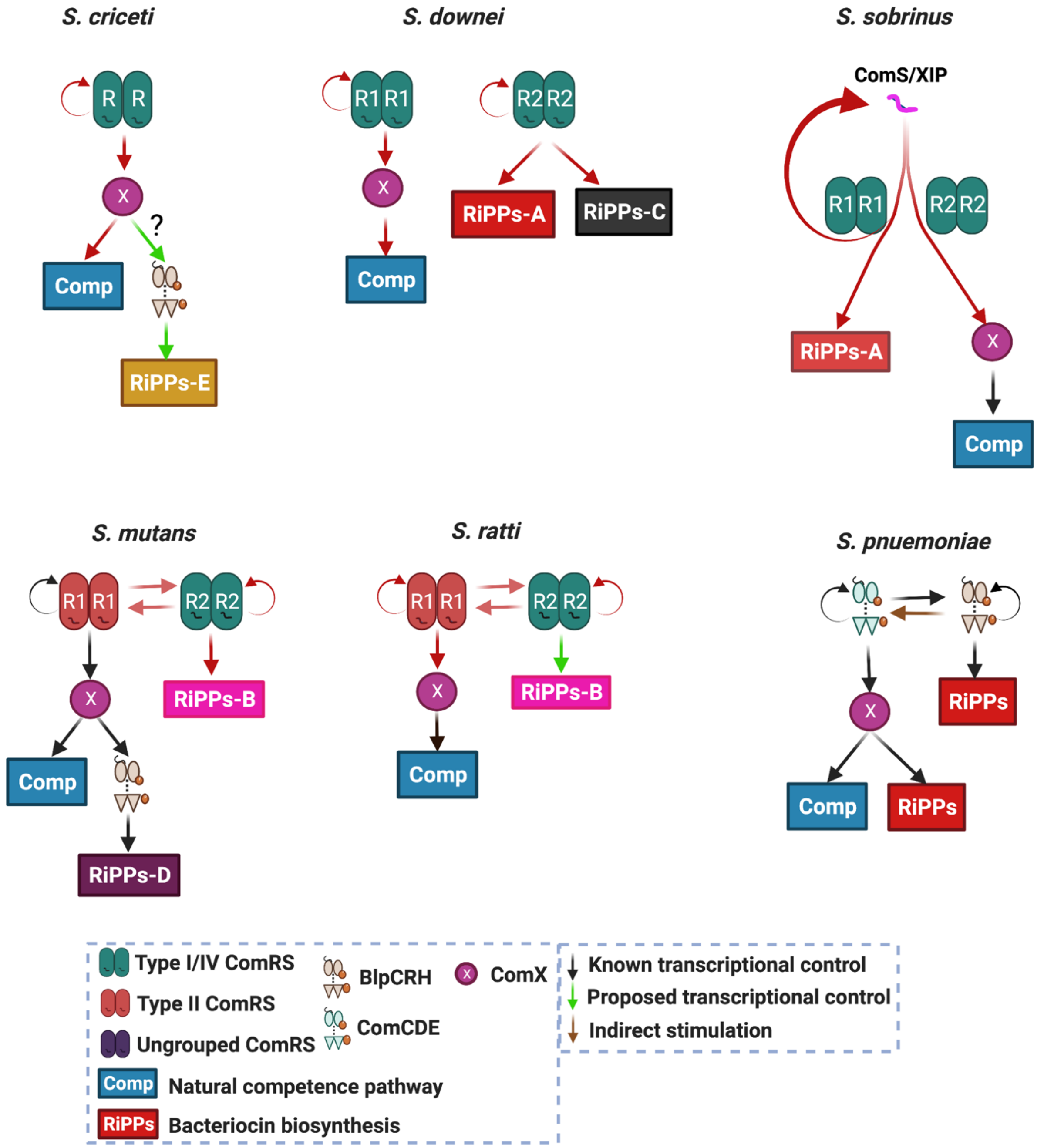

**Supplementary Figure 11. Detailed figures for dissected ComRS networks in mutans streptococci, and comparison with *S. sobrinus* ComCDE-BlpCRH network.**

**References for Supplementary Information:**

[1] J. I. Tietz *et al.*, “A new genome-mining tool redefines the lasso peptide biosynthetic landscape,” *Nat Chem Biol*, vol. 13, no. 5, pp. 470–478, May 2017, doi: 10.1038/nchembio.2319.

[2] M. J. Sullivan, N. K. Petty, and S. A. Beatson, “Easyfig: a genome comparison visualizer,” *Bioinformatics*, vol. 27, no. 7, pp. 1009–1010, Apr. 2011, doi: 10.1093/bioinformatics/btr039.
